# Meiotic pairing through barcode-like satellite DNA repeats

**DOI:** 10.64898/2026.01.06.697918

**Authors:** Lena Skrutl, Ankita Chavan, Anna Sintsova, Ilaria Ceppi, Corin J. Ropp, Shinichi Sunagawa, Petr Cejka, Madhav Jagannathan

## Abstract

During meiosis, chromosomes must find, pair, and synapse with their homologous partners in the crowded milieu of the nucleus^1^. Although homology detection is generally attributed to recombination, pairing and synapsis can occur in its absence^2–10^, suggesting alternate mechanisms that discriminate between homologous and non-homologous chromosomes. In many eukaryotes, tandem repeats known as satellite DNA facilitate inter-chromosomal associations^11^. Notably, their non-uniform distribution across chromosomes gives rise to homologue-specific satellite DNA ‘barcodes’^12–14^, which have been speculated to enable meiotic pairing^15–18^. However, satellite DNA function remains actively debated since these repeats cannot be manipulated in most model organisms. Here, we use satellite DNA deletion, duplication, and translocation strains that are unique to *Drosophila* to demonstrate that repeat mismatches perturb meiotic pairing, particularly at centromeres and pericentromeres. In the absence of satellite DNA homology, pairing is antagonized by the HORMAD protein, Mad2, while a Pachytene checkpoint 2 (Pch2)-dependent meiotic delay restores pairing. Remarkably, pairing defects are also observed in the progeny of *D. melanogaster* natural populations that have diverged in their satellite DNA content. Finally, compromised meiotic pairing is strongly correlated with mid-oogenesis cell death, a quality control mechanism that likely culls defective oocytes to prevent chromosome mis-segregation and aneuploidy. These findings resolve a long-standing debate on satellite DNA functionality by demonstrating a barcode-like role in homology detection. We propose that this repeat-based pairing mechanism exerts an underappreciated selective pressure, constraining the divergence of rapidly evolving satellite DNA within interbreeding natural populations.

## Main

Meiosis, the production of haploid gametes from diploid precursors, is a central feature of sexual reproduction. To ensure the production of gametes with the correct karyotype, it is imperative that the homologous chromosomes segregate to opposite poles during the first meiotic division^1^. This is ensured by a specialized cellular program during meiotic prophase where homologous chromosomes pair along their lengths, synapse and form recombination-dependent physical linkages known as chiasmata^1^. As such, meiotic errors, such as improper pairing and synapsis, can lead to aneuploidy, reproductive difficulties and developmental defects in the offspring^19^.

Pairing requires that homologous chromosomes first find each other in the meiotic nucleus. Across a wide range of species, this is typically achieved by anchoring defined chromosomal loci (e.g. telomeres or centromeres) at the nuclear periphery, effectively reducing the search space for partners from three dimensions to two dimensions^1,20^. Concurrent cytoskeleton-mediated rapid nuclear movements/rotations lead to the clustering of anchored loci, where homologous chromosomes must correctly detect each other, pair and subsequently synapse along their lengths^1^. In many organisms, recombination-dependent homology detection ensures pairing and synapsis with the correct chromosomal partner^1^. However, recombination-independent pairing and synapsis also occurs in many species^2–10,21,22^, hinting at alternate mechanisms that can guide partner choice. These alternate mechanisms may even function in stabilizing homologue associations and safeguarding meiotic fidelity under conditions when chromosomes occasionally fail to be physically linked by recombination, for example during female meiosis in humans^23–27^.

The pericentromeric heterochromatin in many eukaryotes is characterized by the presence of vast tracts of non-coding tandem repeats known as satellite DNA^12^. Typically, each eukaryote contains multiple satellite DNA repeats, which are non-uniformly distributed across the chromosomes^14,28^. While the same satellite DNA repeat can be present on more than one chromosome, each pair of homologous chromosomes contains a unique ‘barcode-like’ array of satellite DNA repeats that is distinct from the other chromosome pairs. In *Drosophila melanogaster*, at least 19 satellite DNA repeats are distributed across the four chromosome pairs^13,28,29^, thus constituting homologue-specific satellite DNA barcodes (Fig. 1a, b). These observations raised the notion that satellite DNA repeats may promote the pairing and synapsis of homologous chromosomes during *Drosophila* meiosis^15–18^. This hypothesis was indirectly supported by the fact that polymerization of the zipper-like synaptonemal complex (SC) largely originates from the centromeric and pericentromeric heterochromatin during zygotene in *Drosophila* oocytes^30,31^. In addition, the pericentromeric heterochromatin of homologous chromosomes remains tightly associated during meiotic prophase in a broad range of species^1,17,30–36^. However, *Drosophila* strains carrying large satellite DNA deletions are not reported to exhibit mutant meiotic phenotypes, such as defective pairing or non-disjunction^12,37^. Thus, the role of satellite DNA in meiosis remains controversial.

**Fig. 1.**
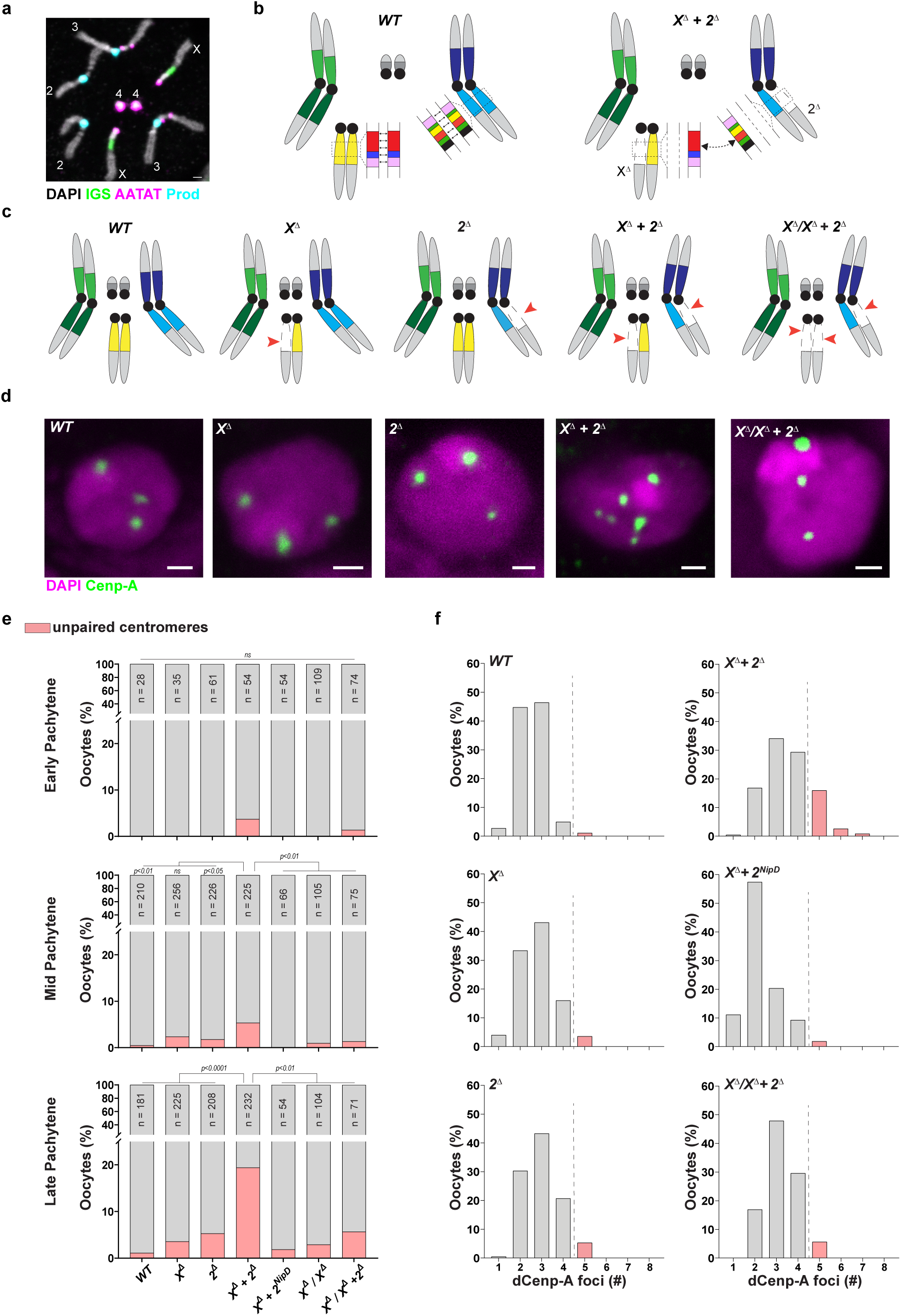
Pericentromeric satellite DNA repeats function in meiotic centromere pairing. **a,** DNA FISH against the IGS satellite repeat (green), the (AATAT)_n_ satellite repeat (magenta) and the *Prod*/(AATAACATAG)_n_ satellite repeat (cyan), co-stained with DAPI (white) on mitotic chromosomes from WT *Drosophila*. **b,** Schematic of the *Drosophila* karyotype with chromosome-specific satellite DNA arrays or satellite DNA ‘barcodes’. Each satellite DNA barcode is made up of individual repeats (red, blue, light pink etc), shown here for two chromosomes, which are capable of self-association. These individual repeats may mediate non-homologous associations (dashed arrow, right panel) when the satellite DNA barcodes are mismatched. **c,** Schematic of the genotypes used in this figure. Red arrows indicate satellite DNA deletions. **d,** IF against dCenp-A (green) in late pachytene oocytes co-stained with DAPI (magenta). **e,** Quantification of oocytes with >4 dCenp-A foci, indicating unpaired centromeres, in early, mid and late pachytene in the indicated genotypes. *n* indicates the number of oocytes analyzed and *p* values were obtained from a Fisher’s exact test. ns indicates *p* > 0.05. **f,** Histogram showing the distribution of number of dCenp-A foci from **(e)**. All scale bars are 1 µm.

### Pericentromeric satellite DNA deletions trigger centromere unpairing during meiotic prophase in *Drosophila*

Meiosis in *Drosophila* ovaries proceeds in a largely canonical manner, with the different prophase stages corresponding to distinct stages of oogenesis (Extended Data Fig. 1a)^38^. In each ovariole, pairing occurs progressively between homologues in pre-meiotic germ cell cysts in region 1 of the germarium^34,35^. During early pachytene, the SC primarily initiates from the 1-2 centromere clusters (Region 2a of the germarium), followed by a slighly later initiation at the chromosomal arms (Region 2b/3). The SC is fully assembled between the paired homologues by mid pachytene (Stg. 2-4 of oogenesis), and is disassembled, albeit incompletely, in late pachytene (Stg. 5-6)^38^ (Extended Data Fig. 1a). The relatively simple karyotype of *D. melanogaster* (2n=8) means that the presence of >4 centromeric foci in oocytes, labelled by a dCenp-A antibody, unambiguously indicates unpaired centromeres. Importantly, 5 or more dCenp-A foci are observed in oocytes where components of the chromosome axis or the SC are depleted^30,31^. Therefore, quantification of centromere foci in oocytes offers a straightforward readout for meiotic pairing.

We tested the role of satellite DNA repeats in meiotic pairing using two well-characterized satellite DNA deletion strains. First, we used the *Zhr^1^* allele (hereafter *X^Δ^*), an X;Y compound chromosome, where the long arm of the X chromosome, which is missing nearly all of the ∼11Mb pericentromeric 359bp satellite DNA repeat^13,39,40^, is fused to a small fragment from the Y chromosome (Fig. 1c, Extended Data Fig. 1b). We also used the *M41A10* allele (hereafter *2^ý^*), which is missing nearly all the satellite DNA repeats and 41 heterochromatin-embedded coding genes on Chr. 2R^13,41^ (Fig. 1c, Extended Data Fig. 1c). Since these satellite DNA deletions can potentially affect the expression of heterochromatin-embedded or heterochromatin-proximal genes, all experiments (unless otherwise specified) are performed in strains that are heterozygous for these deletions (*X^Δ^*/*+* or *2^Δ^*/*+*), hereafter simply referred to as *X^Δ^* and *2^Δ^* (Fig. 1c). Despite the structural heterozygosity between the satellite DNA deletion chromosomes and their wildtype homologues, centromere pairing was largely intact in the satellite DNA deletion strains (*X^Δ^*or *2^Δ^*) in pachytene oocytes and comparable to the wildtype (Fig. 1d-f). These data, as well as similar observations from the past^37^, have largely driven the conclusion that satellite DNA repeats are dispensable for meiotic pairing.

However, we reasoned that the loss of satellite DNA repeats from a single chromosome would likely only hinder the association of the affected homologue pair, while the association between all other intact homologue pairs would be unaffected. We therefore predicted that the chromosome carrying the satellite DNA deletion, due to a lack of alternative options, would eventually associate with its correct partner, likely through remnant satellite DNA repeats, recombination or other redundant homology detection pathways^42–45^. We considered that this ‘pairing by default’ may explain the previously observed lack of pairing defects in strains carrying satellite DNA deletions on a single chromosome. Therefore, we hypothesized that satellite DNA deletions from at least two non-homologous chromosomes would be required to assess whether satellite DNA repeats function in meiotic pairing (Fig. 1b, right panel). To test our hypothesis, we quantified centromere pairing in strains heterozygous for both *X^Δ^*and *2^Δ^* (hereafter ‘*X^Δ^*/+; *2^Δ^*/+’ or ‘double deletion’). While only a modest increase in centromere unpairing was detected in early and mid pachytene, we observed that 19.4% of late pachytene oocytes from double deletion strain exhibited >4 centromeric foci, a ∼4-fold increase in comparison to the controls (Fig. 1d-f). The reason for defective centromere pairing being mainly observed in late pachytene is explained below. Nevertheless, our data are a clear indication that deletion of satellite DNA repeats can compromise meiotic pairing.

In addition to the loss of satellite DNA repeats, the *2^Δ^* chromosome also lacks 41 genes at the Chr. 2R heterochromatin boundary, including essential genes such as *RpL38* and genes important for chromosome structure and maintenance such as the Tip60 complex subunit, *Nipped-A*, and the cohesin loader, *Nipped-B*^46–48^. Given the importance of pericentromeric cohesion for meiotic pairing, we considered that pairing phenotypes in the double deletion strain could be due to a genetic interaction between the deleted satellite DNA on the *X^Δ^* chromosome and gene heterozygosity on the *2^Δ^* chromosome. To control for this, we used a Chr. 2R deletion allele, *Df(2R)^Nip-D^* (hereafter *2^NipD^*)^49^, which lacks *RpL38*, *Nipped-A*, *Nipped-B* and 18 other genes that are deleted in 2^Δ^ but does not extend into the pericentromeric satellite DNA (Extended Data Fig. 1d). In contrast to the *X^Δ^*; *2^Δ^* strain, we observed that flies heterozygous for both *X^Δ^* and *2^NipD^* did not exhibit centromere unpairing (Fig. 1e, f, Extended Data Fig. 1e, f). We also used transcriptome-wide gene expression analysis to determine whether Chr. 2R heterochromatin boundary genes were misexpressed in ovaries from the double deletion strain in comparison to ovaries from the other genotypes. Although the whole ovary transcriptome of each of the genotypes was distinct (Extended Data Fig. 2a), none of the Chr. 2R heterochromatin proximal genes were specifically misexpressed (log_2_FC>1, p_adj_<0.05) in the double deletion strain in comparison to all the other genotypes (Extended Data Fig. 2b). Thus, we conclude that the meiotic pairing defects in the double deletion strain are not caused by misexpression of Chr. 2R heterochromatin-proximal genes.

### Satellite DNA deletion chromosomes are sensitive to pairing defects

Since non-homologous centromeres remain clustered across meiotic prophase in *Drosophila*^38^, we considered that our quantification of centromere foci may underestimate unpairing. Therefore, we used probes against the Responder (Rsp) and Dodeca satellite DNA repeats, which are uniquely present at the pericentromeric heterochromatin of Chr. 2 and Chr. 3 respectively^28^, to directly assess pairing at specific chromosomes. We observed that the pericentromeric heterochromatin of Chr. 2 (Rsp satellite DNA repeat) was unpaired in 28.2% of late pachytene double deletion oocytes (Fig. 2a-c), a >2-fold increase in comparison to the controls. In contrast, the pericentromeric heterochromatin of Chr. 3 (Dodeca satellite DNA repeat) remained paired across all genotypes and all pachytene stages (Fig. 2a, b, d). Notably, the Rsp locus, but not Dodeca, was also unpaired in earlier stages of pachytene (Fig. 2a-c), as well as in oocyte precursors in region 1 of the germarium (Fig. 2a-c, Extended Data Fig. 1a), where homologue pairing is initially established^34,35^. These data highlight that homologue-specific unpairing can occur, even in oocytes with fewer than 5 dCenp-A foci. While quantification of centromere foci likely underestimates the full extent of unpairing, we note that this mainly affects the double deletion strain. Overall, our data indicate that chromosomes lacking their full array of satellite DNA repeats are sensitive to pairing defects.

**Fig. 2.**
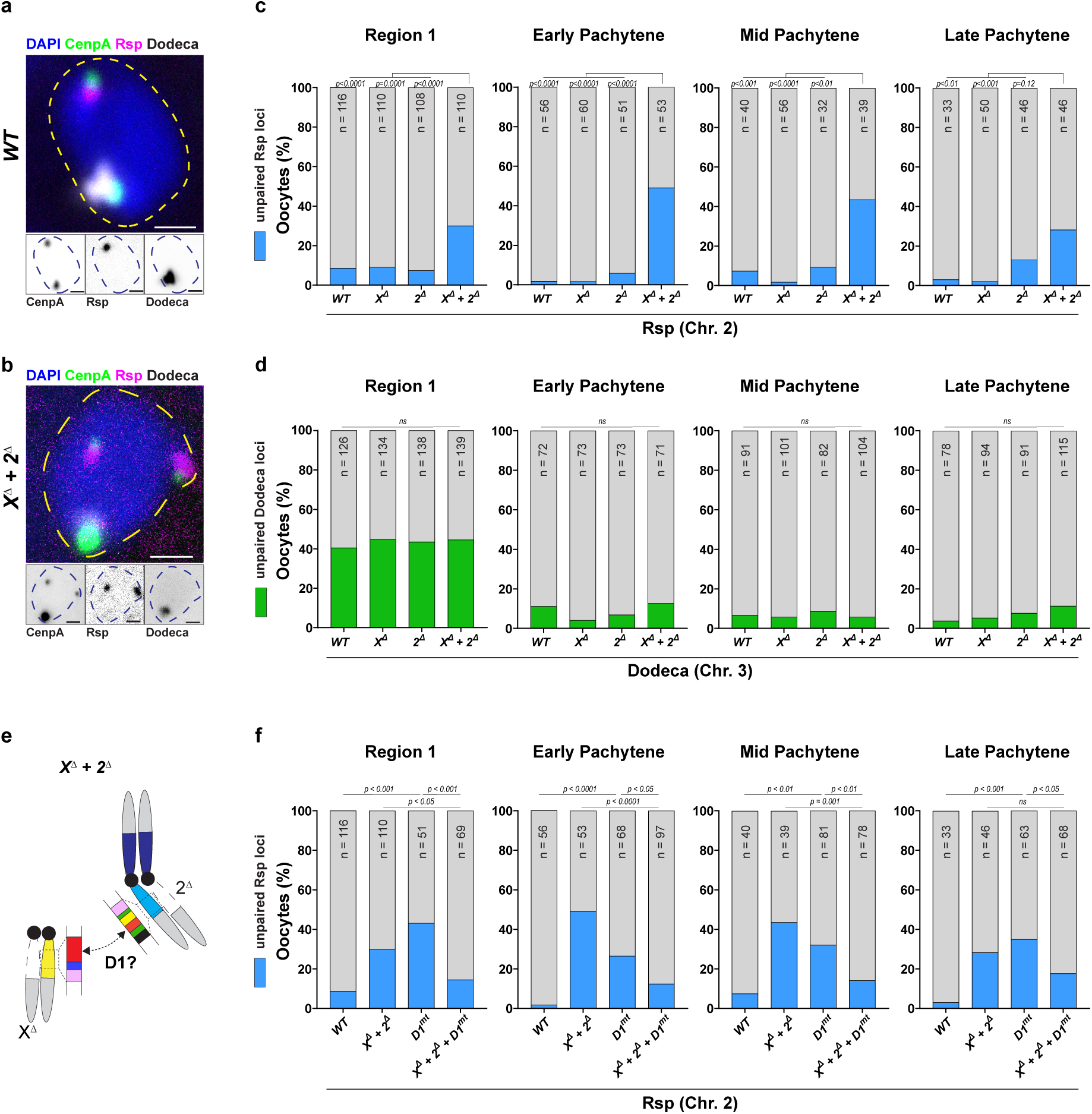
Loss of meiotic centromere pairing at chromosomes carrying satellite DNA deletions. **a, b,** DNA FISH against the Chr. 2 specific Rsp satellite repeat (magenta) and the Dodeca satellite repeat (white) in mid pachytene oocytes from the WT strain (a) and the double deletion strain (b) and co-stained with DAPI (blue) and dCenp-A (green). **c,d,** Quantification of oocytes with unpaired Rsp (c) and Dodeca (d) loci in region 1, early, mid, and late pachytene from the indicated genotypes. **e**, Schematic showing non-homologous associations between identical satellite DNA repeats (red) in the double-deletion strain, potentially mediated by the satellite DNA-binding protein D1. **f**, Quantification of oocytes with unpaired Rsp loci in region 1, early, mid, and late pachytene from the indicated genotypes. *n* indicates the number of oocytes analyzed and *p* values were obtained from a Fisher’s exact test. ns indicates *p* > 0.05. All scale bars are 1 µm.

### Pairing defects mainly occur at centromeric and pericentromeric heterochromatin

Despite the occurrence of centromere and pericentromere unpairing in the double deletion strain, synaptonemal complex (SC) morphology during pachytene was surprsingly consistent between all genotypes (Extended Data Fig. 3a). For a higher-resolution snapshot of pairing at the chromosome arms, we used Oligopaint-based DNA FISH^50^ at three ∼1Mb loci on Chr. 2R (Extended Data Fig. 3b). In *WT* oocytes, these loci were largely paired during mid pachytene and unpaired in late pachytene, when the SC begins to be disassembled (Extended Data Fig. 3c-g). These three loci were also paired during mid pachytene in all the satellite DNA deletion genotypes (Extended Data Fig. 3c-g), indicating that the meiotic pairing defects in the double deletion strain are largely confined to centromeres and pericentromeres.

### Satellite DNA associations between non-homologous chromosomes likely underlie centromere unpairing

Our ‘pairing by default’ model predicts that the homozygous deletion of satellite DNA repeats will not lead to pairing defects, since all other intact chromosomes remain properly associated and the satellite DNA-deleted pair will eventually end up with each other, likely through redundant homology detection pathways. Consistently, we did not observe centromere pairing defects in a strain homozygous for *X^Δ^* (Fig. 1e, Extended Data Fig. 1g, h). In addition, these data indicate that the total amount of deleted satellite DNA (similar between *X^Δ^* homozygous strain and the double deletion strain) is not the underlying cause of centromere unpairing. Rather, we hypothesized that non-homologous satellite DNA associations between the WT Chr. X and the WT Chr. 2 in the double deletion strain may drive centromere unpairing (dashed arrow, right panel, Fig. 1b). To test this, we first generated a strain homozygous for *X^Δ^* and heterozygous for *2^Δ^* (*X^Δ^*/ *X^Δ^*; *2^Δ^*/+) (Fig. 1c) and found that centromere pairing was restored across pachytene, in comparison to the double deletion (*X^Δ^*/+; *2^Δ^*/+) strain (Fig. 1d-f). Our data suggest that under conditions where the entire satellite DNA array is not fully engaged with its homologue, interactions between similar or identical satellite DNA repeats on non-homologous chromosomes may impair homologue pairing (Fig. 1b). Thus, the same repeats, which are important for pairing, may also hinder the process if the satellite DNA barcode is disrupted.

### The D1 satellite DNA-binding protein mediates non-homologous associations in the double deletion strain

We next sought to better understand the non-homologous satellite DNA associations, likely between the intact Chr. X and Chr. 2, that underlie the pairing defects in the double deletion strain. The *X^Δ^* allele lacks ∼11Mb of the pericentromeric AT-rich 359bp satellite DNA repeat (Extended Data Fig. 4a) and the rescue of centromere pairing in the *X^Δ^*/ *X^Δ^*; *2^Δ^*/+ strain suggests that unengaged 359bp repeats on *X^WT^* may be required for the non-homologous associations. A potential target on Chr. 2 is the 359bp-derived 260bp satellite DNA repeat^51,52^. Interestingly, an AT-hook containing satellite DNA-binding protein known as D1 localizes to the X chromosome pericentromeric heterochromatin and 359bp-containing nucleosomes^53,54^, and has been demonstrated to mediate inter-chromosomal satellite DNA associations^55,56^. Using an electrophoretic mobility shift assay (EMSA), we found that purified D1 specifically and directly binds both the 359bp and 260bp repeat monomers, in comparison to 50% GC-containing control sequences of the same length (Extended Data Fig. 4b-c). We also found that the addition of D1 to micron-sized paramagnetic beads coated with DNA (50%GC control, 260bp satellite DNA monomer, 359bp satellite DNA monomer) was sufficient to induce bead clustering, in a manner that scaled with DNA-binding affinity (Extended Data Fig. 4d-f). In vivo, we observed that D1 co-localizes with the 359bp repeat in female germ cells (Extended Data Fig. 4g). Notably, the 260bp repeat, as well as another D1-bound satellite DNA repeat (AATAG)^56^, are in excess on the *2^WT^* chromosome, in comparison to *2^Δ^*(Extended Data Fig. 4h). These observations led us to hypothesize that D1-mediated satellite DNA associations may be responsible for centromere unpairing in the double deletion strain (Fig. 2e). To do so, we introduced a CRISPR-generated D1 knockout allele (*D1^KO^*)^56^ into the double deletion background (*X^Δ^*/+; *2^Δ^*/+; *D1^KO^*/*D1^KO^*) and assessed pairing at the Rsp locus. We observed that loss of D1 in the double deletion background significantly rescued pairing at Chr. 2 pericentromeric heterochromatin across region 1 as well as early and mid pachytene (Fig. 2f). We note that loss of D1 alone resulted in unpairing at the Chr. 2 pericentromeric heterochromatin, which we attribute to a pairing function for D1 in a *WT* background, likely on bound satellite DNA repeats such as AATAG and 260bp. We propose that the same function of D1 mediates non-homologue associations when D1-bound repeats (e.g. 359bp, 260bp and AATAG) are unengaged. Overall, these data indicate that D1-dependent associations between bound satellite DNA repeats on non-homologous chromosomes drives unpairing in double deletion oocytes.

### Associations between the chromosome arms promotes centromere pairing in the absence of satellite DNA repeats

We were intrigued by the fact that double deletion oocytes only exhibited >4 dCenp-A foci in late pachytene (Fig. 1e, f), despite clearly showing pairing defects at the pericentromeric heterochromatin in earlier stages (Fig. 2c). One reason for this discrepancy could be the phenomenon of centromere clustering^30,31,35^, which likely masks the full extent of homologue unpairing. However, we also considered that synapsis and recombination at the chromosomal arms, which can reorient incorrectly paired homologues^57^, may partly explain the lack of >4 dCenp-A foci in early and mid pachytene oocytes from the double deletion strain. Following this line of reasoning, the full extent of centromere unpairing would only become apparent upon SC disassembly in late pachytene oocytes, which is consistent with our data (Fig. 1e). Therefore, we sought to assess centromere unpairing in mid pachytene oocytes from the double deletion strain lacking homologue synapsis or recombination. Since loss of SC components are known to generally affect centromere clustering and pairing in *Drosophila*^30,31^, we instead used a 2^nd^ chromosome balancer (hereafter *CyO*), which likely weakens synapsis with a *WT* 2^nd^ chromosome due to multiple inversions at the chromosomal arms (Extended Data Fig. 5a)^58,59^. We observed significantly higher levels of centromere unpairing across early, mid and late pachytene in the *X^Δ^*/+; *2^Δ^*/*CyO* strain, in comparison to the single deletion control strain (*X^Δ^*/+; +/*CyO*) as well as the double deletion strain (*X^Δ^*/+; *2^Δ^*/+) (Extended Data Fig. 5b, c). To test the role of recombination, we used a validated loss-of-function allele of *mei-P22* (*mei-P22^P22^*)^2^, which is essential for the induction of meiotic DSBs by the *Drosophila spo11* orthologue, *mei-W68*. Consistent with our prediction, we found that introducing the *mei-P22^P22^* mutation into the double deletion background led to a >5-fold increase in centromere unpairing across early and mid pachytene, in comparison to the double deletion oocytes (Extended Data Fig. 5b, c). Thus, synapsis and recombination at the chromosomal arms partially suppress the centromere unpairing phenotype in the double deletion strain.

### Mismatches in chromosome-specific satellite DNA barcodes elicit centromere unpairing

We next asked whether changing the composition of the satellite DNA ‘barcodes’ could have similar effects on meiotic pairing. We first used a strain (*Df(2L)c’*, hereafter *2^Dup^*), where a large fraction of the Chr. 2R satellite DNA repeats are duplicated onto the pericentromeric heterochromatin of Chr. 2L (Fig. 3a), thus disrupting the *WT* arrangement of satellite DNA repeats^41^. In strains heterozygous for *2^Dup^*, we did not observe centromere unpairing, likely because pairing between all other homologous chromosomes was unaffected (Fig. 3b-d). However, introducing the *2^Dup^* chromosome into the *X^Δ^* background (*X^Δ^*/+; *2^Dup^*/+) was sufficient to induce centromere unpairing across mid and late pachytene (Fig. 3b-d). This pairing defect was further exacerbated when *2^Dup^* was introduced into the double deletion background (*X^Δ^*/+; *2^Δ^*/*2^Dup^*, Fig. 3b-d). Based on these data, we propose that the mismatched satellite DNA repeats on *2^Dup^*, when given the option, associate with similar repeats on non-homologous chromosomes, thus driving centromere unpairing.

**Fig. 3.**
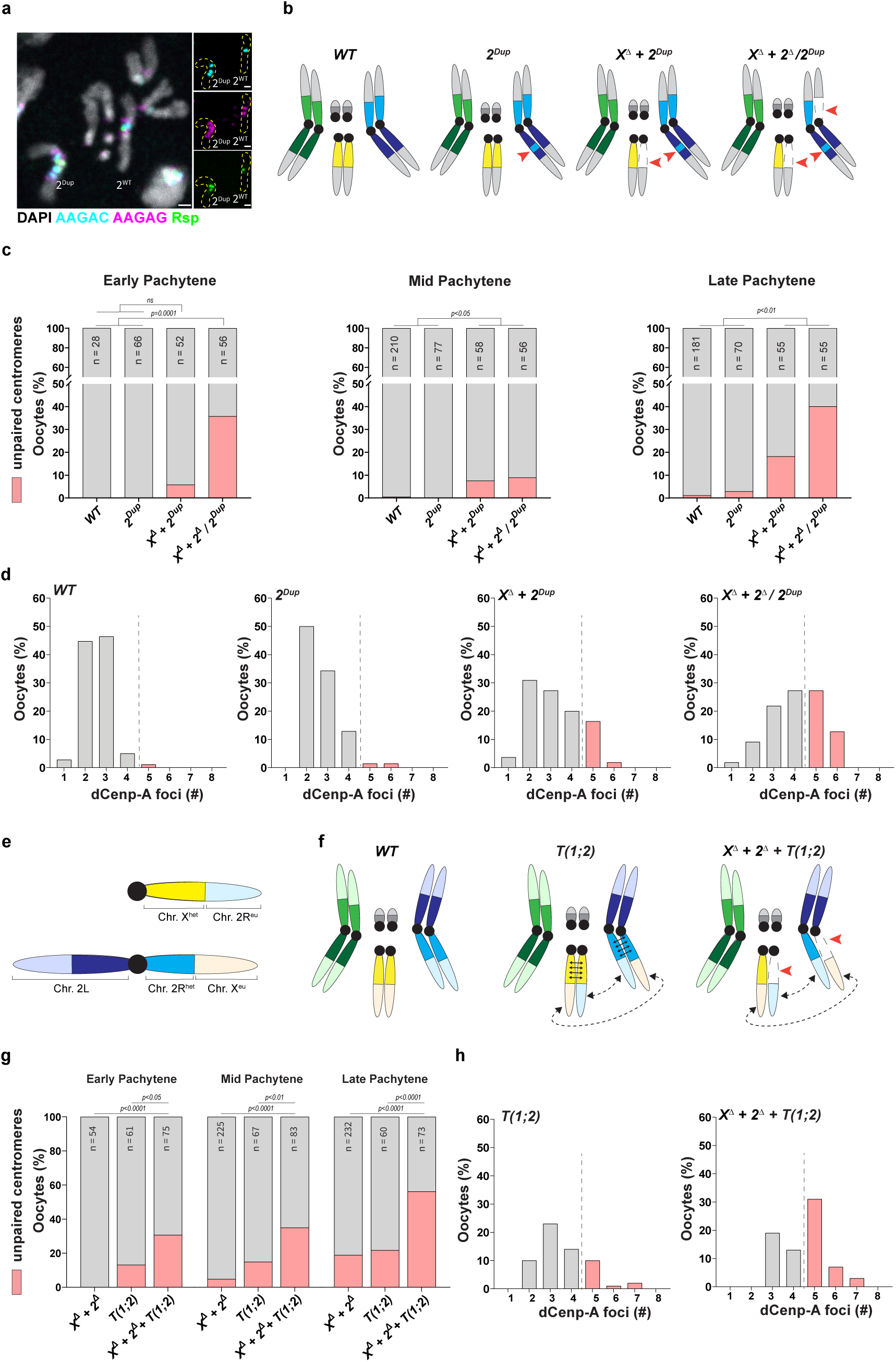
Mismatched satellite DNA repeats drive meiotic centromere unpairing. **a,** DNA FISH against the Rsp satellite repeat (green), the (AAGAG)_n_ satellite repeat (magenta) and the (AAGAC)_n_ satellite repeat (cyan) and co-stained with DAPI (white) on mitotic chromosomes from a strain heterozygous for *2^Dup^*. **b,** Schematic of the genotypes used in this figure. Each colour represents a chromosome arm-specific satellite DNA barcode. Red arrows indicate satellite DNA deletions or duplications. **c,** Quantification of oocytes with >4 dCenp-A foci, indicating unpaired centromeres, in early, mid and late pachytene in the indicated genotypes. **d,** Histogram showing the distribution of number of dCenp-A foci from **(c)**. **e,** Schematic of the T(1;2) translocation strain with heterochromatin-proximal breakpoints at cytological bands 19-20 on Chr. X and cytological bands 40-41 on Chr. 2. Bright colors indicate the heterochromatic satellite DNA barcodes and the fainter colors indicate the corresponding euchromatic arms. **f,** Schematic of the translocation genotypes used in **(g,h)**. Red arrows indicate satellite DNA deletions. Solid lines represent association between homologous satellite DNA barcodes while dashed lines indicate synapsis-dependent association between the translocated chromosomal arms. **g,** Quantification of oocytes with >4 dCenp-A foci, indicating unpaired centromeres, in early, mid and late pachytene in the indicated genotypes. *n* indicates the number of oocytes analyzed and *p* values were obtained from a Fisher’s exact test. ns indicates *p* > 0.05. **h,** Histogram showing the distribution of number of dCenp-A foci from **(g)**. WT and *X^ý^* + *2^ý^* unpaired centromere values were reused from Fig. 1e.

To determine the extent to which satellite DNA repeats play an instructive role in meiotic pairing, we utilized a X;2 translocation strain, *T(1;2)*, where the breakpoints occur near the euchromatin-heterochromatin boundary on Chr. X and Chr. 2R (Fig. 3e). As a result, the right arm of Chr. 2 is fused to the X chromosome pericentromeric heterochromatin, while the euchromatic arm of the X chromosome is fused to the pericentromeric heterochromatin of Chr. 2R (Fig. 3e). In the *T(1;2)* strain, we observed centromere unpairing at levels comparable to the double deletion strain (Fig. 3f-h). We considered that this defect could be driven by pairing between the translocated arms, which would oppose the pairing between homologous satellite DNA arrays. In this model, further weakening satellite DNA associations should exacerbate the pairing defect. To test this, we introduced the *T(1;2)* translocation into the double deletion background (Fig. 3f) and observed a stronger effect on centromere unpairing across early, mid and late pachytene in comparison to both the double deletion and *T(1;2)* strains (Fig. 3f-h). Therefore, our data clearly indicate that homologous satellite DNA arrays are paired under normal circumstances, irrespective of the composition of the rest of the chromosome.

### Satellite DNA variation in natural populations is associated with pairing defects

A five-continent reference panel consisting of 84 *D. melanogaster* wild strains^60^, termed the ‘Global Diversity Lines (GDLs)’ allowed us to test whether variation of satellite DNA repeats in natural populations is associated with meiotic unpairing (Fig. 4a). In-depth sequencing of the GDL collection has characterized the single nucleotide polymorphisms (SNPs), indels and satellite DNA profiles of the individual strains^60,61^. We performed a phylogenetic analysis of ‘neutral’ SNPs (SNPs from small introns and fourfold degenerate coding positions^60^), and found, as expected, that the GDL strains clustered by geographical origin (Extended Data Fig. 6a, b). Similarly, strains sharing the same geographical origin also clustered together based on indels (Extended Data Fig. 6c). In contrast, a principal component analysis (PCA) using previous estimates of GDL strain satellite DNA content^61^ did not reveal a geographical relationship (Fig. 4b). Rather, strains from different geographical regions clustered together while strains from the same geographical region exhibited significant satellite DNA variation (Fig. 4b), consistent with the rapid evolution of satellite DNA repeats. We used the satellite DNA PCA to select pairs of GDL strains containing similar satellite DNA content (*N 02* x *N 14* and *I 38* x *B 52*) and containing divergent satellite DNA content (*I 16* x *I 22* and *I 16* and *ZH 33*) and assessed centromere pairing in the progeny of these strain pairs in late pachytene. We found that the progeny of strain pairs with divergent satellite DNA content exhibited significant levels of centromere unpairing, in contrast to the progeny of strain pairs with similar satellite DNA content (Fig. 4c). To ensure that the strains (*I16*, *I22* and *ZH33*) whose progeny exhibited centromere unpairing were not inherently defective, we crossed each of them to another GDL strain with a similar satellite DNA profile (Fig. 4b). Importantly, we found that the progeny of these GDL pairs (*I16* x *B59*, *I22* x *T22A* and *ZH33* x *B42*) did not exhibit centromere unpairing (Fig. 4c). Moreover, we observed a strong correlation between satellite DNA divergence and centromere unpairing (Extended Data Fig. 6d), while no relationship was observed between SNP- and indel-based divergence and centromere unpairing (Extended Data Fig. 6e, f). Overall, our data suggest that pairing defects are largely driven by differences in satellite DNA content, and not other variation intrinsic to these strains.

**Fig. 4:**
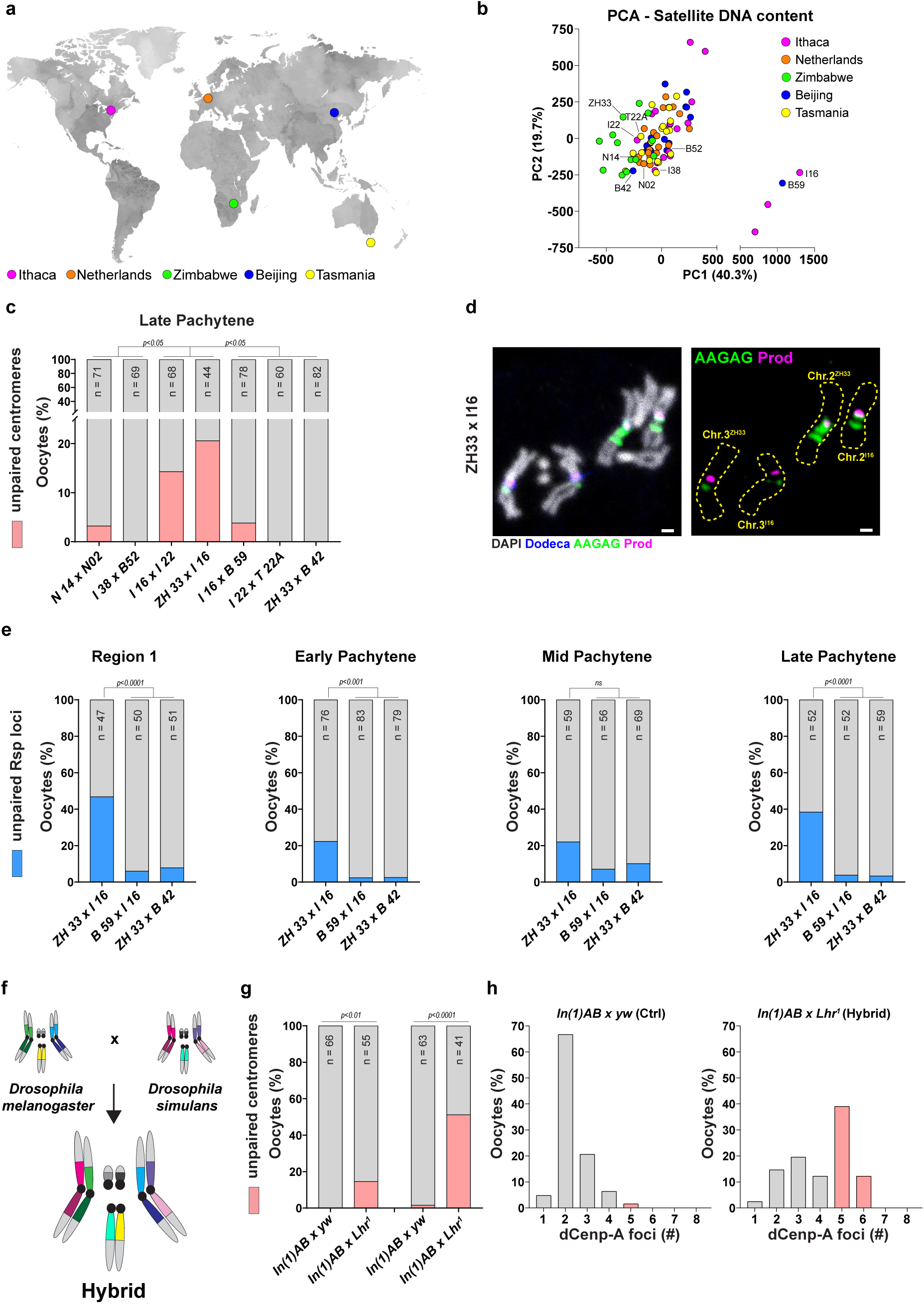
Satellite DNA divergence in naturally occurring populations leads to meiotic pairing defects. **a,** Map indicating the geographical origin of strains from the Global Diversity Lines (GDL) collection. **b,** Principal component analysis (PCA) of satellite DNA content from the GDL strains based on previous estimates^61^. **c,** Quantification of oocytes with >4 dCenp-A foci, indicating unpaired centromeres, in late pachytene in the indicated genotypes. **d**, FISH against the Dodeca satellite repeat (blue), (AAGAG)_n_ satellite repeat (green), Prod/(AATAACATAG)_n_ satellite repeat (magenta) co-stained with DAPI (white) on mitotic chromosomes in the progeny of ZH33 and I16 strains. **e,** Quantification of oocytes with unpaired Rsp loci in Region 1, early, mid, and late pachytene from the indicated genotypes. **f,** Schematic of satellite DNA barcode mismatches in inter-species hybrids. **g,** Quantification of oocytes with >4 dCenp-A foci, indicating unpaired centromeres, in mid (left) and late pachytene (right) in the indicated genotypes. **h,** Histogram showing the distribution of number of dCenp-A foci in late pachytene oocytes from the hybrid and the pure species control from **(g)**. *n* indicates the number of oocytes analyzed and *p* values were obtained from a Fisher’s exact test. ns indicates *p* > 0.05.

We noted that much of the satellite DNA variation in the GDL strains was driven by loss of the AAGAG repeat, which is normally the most abundant satellite DNA repeat in *Drosophila*^13^. This repeat was barely detected in both *I16* and *B59* (Extended Data Fig. 6g). To validate these differences, we directly assessed satellite DNA abundance using DNA FISH on mitotic chromosome spreads in the progeny of *ZH33* x *I16* and *B59* x *I16* (Extended Data Fig. 7a, see methods for details). In the progeny of *ZH33* x *I16*, we observed that the homologue that contained reduced levels of AAGAG (Fig. 4d, Extended Data Fig. 7a, c), exhibited a consistent change in the abundance of other satellite DNA repeats (Fig. 4d, Extended Data Fig. 7d-f). These data suggest altered satellite DNA composition in the chromosomes with reduced AAGAG, likely corresponding to the *I16* homologue. In contrast, a similar change in the satellite DNA barcodes was not observed in the progeny of *B59* x *I16*, where both parental strains are estimated to have similar satellite DNA content (Extended Data Fig. 7b, g-j). Since the majority of AAGAG (∼5.5Mb)^13^ is present on Chr. 2, we assessed pairing at the Rsp locus in the progeny of *ZH33* x *I16*, as well as in the progeny of *ZH33 x B42* and *I16 x B59* (our negative controls). Importantly, we observed a substantial pairing defect in the progeny of *ZH33* x *I16,* including in region 1 oocyte precursors where pairing is established (Fig. 4e). A similar pairing defect was not observed in our negative controls (Fig. 4e), whose parental strains have similar satellite DNA profiles. Overall, our data strongly suggest that variation in satellite DNA content is sufficient to impair the establishment of meiotic pairing in natural populations.

Finally, we took advantage of the vastly different satellite DNA content between *Drosophila melanogaster* and its closest sibling species, *Drosophila simulans*, even though the rest of their genomes retain 85-95% sequence identity^28,62^. While hybrid progeny are agametic with a near complete loss of germ cells, certain strains permit oogenesis in hybrids, upto to the production of mature, albeit inviable, eggs^63,64^. Using these strains, we were able to generate hybrid females, where the satellite DNA barcodes on all chromosomes were markedly different (Fig. 4f). We observed that 14.5% and 51.2% of mid and late pachytene oocytes were unpaired in these female hybrids in comparison to 0.0-1.6% in the pure species control (Fig. 4g, h). Female hybrids also exhibited synapsis defects in mid pachytene, including SC discontinuities and the occurrence of polycomplexes (Extended Data Fig. 7k). Similar SC discontinuities and aberrations were also observed in other genotypes with comparable levels of unpairing (Extended Data Fig. 7k). These data suggest that a persistent inability to pair homologous chromosomes may lead to to irreversible defects in SC polymerization. Taken together, our data implicate satellite DNA repeats as key factors that contribute to meiotic pairing between homologous chromosomes.

### A conserved meiotic checkpoint modulates centromere unpairing in the response to satellite DNA deletions

Synapsis defects in many model organisms triggers a conserved meiotic checkpoint, mediated by a AAA+ ATPase known as Pachytene checkpoint 2 (Pch2, TRIP13 in vertebrates)^65–69^. In *Drosophila*, previous studies have demonstrated that defects in homologue alignment delay oocyte specification in a Pch2-depenent manner during early to-mid pachytene, resulting in region 3 egg chambers containing two SC-positive pro-oocytes (the two-oocyte phenotype, Fig. 5a)^68^. In the double deletion strain, we observed that almost all region 3 egg chambers exhibited the two-oocyte phenotype (Fig. 5a, b), suggesting that the Pch2-dependent meiotic delay was activated. We first mutated *pch2* using validated loss-of-function alleles^70^ and found that centromere pairing was mildly perturbed. These data were reminiscent of *TRIP13^mt^* mouse spermatocytes and oocytes, which also exhibit asynapsis at the satellite DNA-rich centromeric and pericentromeric regions^71^. Next, we mutated *pch2* in the double deletion background and observed that the *X^Δ^*/+; *2^Δ^*/+*; pch2^mt^* strain exhibited centromere unpairing in 31.6 and 37.7% of mid and late pachytene oocytes respectively, a ∼2-4-fold increase in comparison to both the *pch2* mutant and double deletion strains (Fig. 5c). These data suggest that Pch2 activation allows for correct homologue pairing in the absence of full satellite DNA homology, likely through modulating progression through pachytene.

**Fig. 5.**
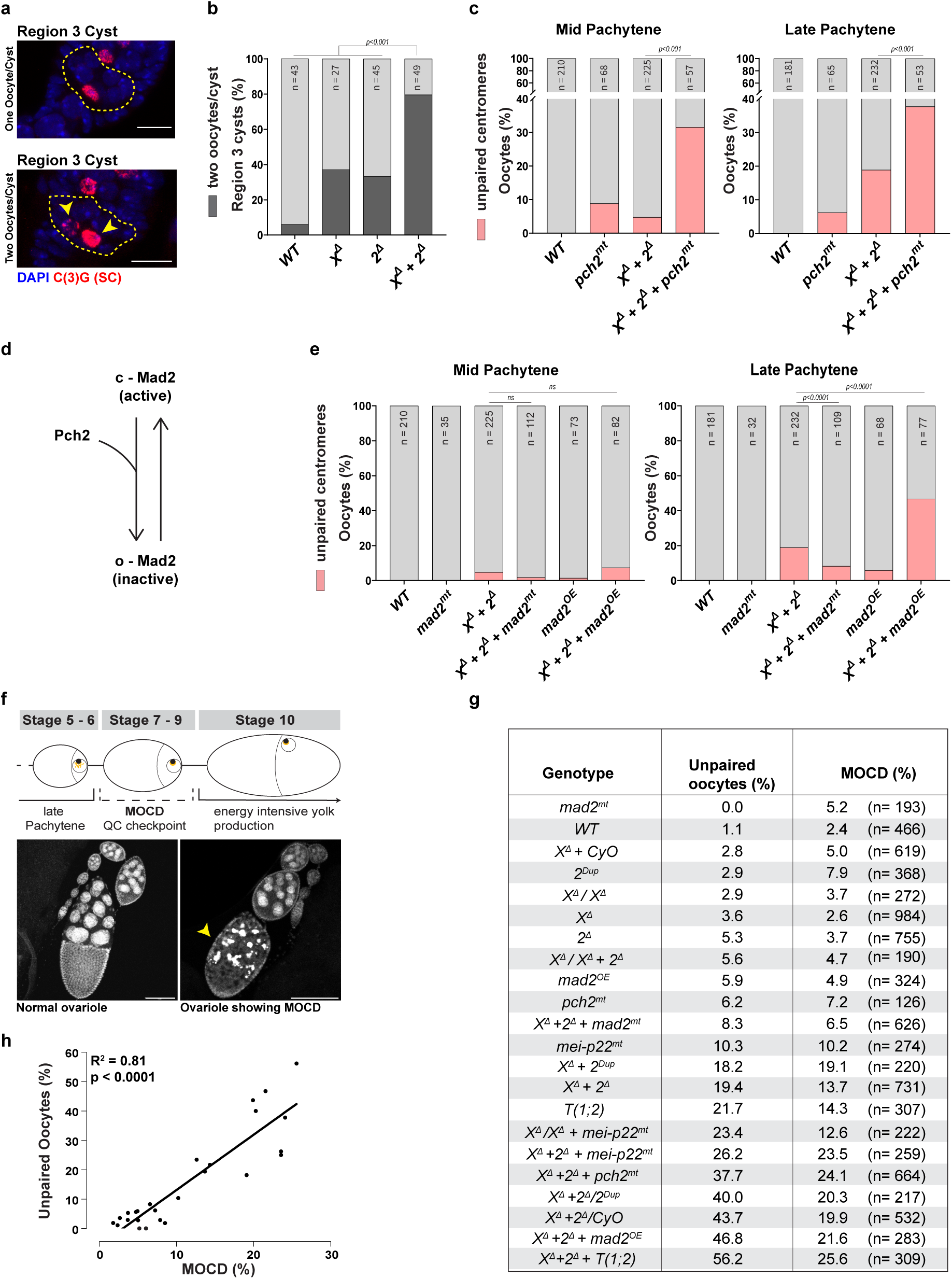
Loss of satellite DNA homology activates meiotic checkpoints and triggers cell death. **a,** IF against the transverse filament of the synaptonemal complex C(3)G (red) in the outlined region 3 cysts and co-stained with DAPI (blue). Arrow indicates the second oocyte with persistent C(3)G signal. **b,** Quantification of region 3 cysts showing the double oocyte phenotype in the indicated genotypes. *n* indicates the number of cysts analyzed and *P* was obtained from a Fisher’s exact test. **c,** Quantification of oocytes with >4 dCenp-A foci, indicating unpaired centromeres, in mid and late pachytene in the indicated genotypes. *pch2* was mutated using the *pch2^EY01788^* allele in trans to *Df(3R)^Bsc195^*. **d,** Schematic showing regulation of Mad2 activity by Pch2. **e,** Quantification of oocytes with >4 dCenp-A foci, indicating unpaired centromeres, in mid and late pachytene in the indicated genotypes. *mad2* was mutated using the *mad2^EY21687^* allele in trans to *mad2^P^*. *mad2^OE^* represents an extra copy of a *GFP::Mad2* transgene. **f,** Schematic of the quality control mid-oogenesis checkpoint (MOCD) at Stg 7-9 of oogenesis. A healthy ovariole (bottom left) and an ovariole exhibiting MOCD (bottom right) are co-stained with DAPI. Arrowhead points to the Stg. 8 egg chamber undergoing MOCD with pyknotic nuclei **g,** Table indicating percentage of oocytes with unpaired centromeres in late pachytene and corresponding percentage of Stg 7-9 egg chambers with MOCD from the indicated genotypes. n indicates total number of Stg 7-9 egg chambers scored. **h**, Correlation between centromere unpairing and MOCD from **(g)**. R^2^ and slope were determined from a linear regression analysis. *n* indicates the number of oocytes analyzed and *p* values were obtained from a Fisher’s exact test. ns indicates *p* > 0.05. WT and *X^ý^* + *2^ý^* unpaired centromere values were reused from Fig. 1e.

Pch2’s best-characterized function involves remodeling proteins that contain a HORMA domain^72^. During meiosis, conserved HORMA-domain containing proteins such as Hop1 in *S. cerevisiae* and *HIM-3* in *C. elegans*, which localize to the chromosomal axis, are thought to be important targets of Pch2^65,73^. However, these meiotic HORMADs are not present in the *Drosophila* genome^65,72^. Therefore, we turned our attention to another HORMAD, Mad2, which mainly functions in the tension-sensing spindle assembly checkpoint (SAC)^72^ during chromosome segregation. Interestingly, Mad2 has also been shown to function during meiotic prophase in *C. elegans*, where it negatively regulates chromosome synapsis^74,75^. Moreover, Pch2 is known to remodel Mad2 from a closed (and active) conformation to an open (and inactive) conformation (Fig. 5d)^65,72^. Therefore, we introduced loss-of-function *mad2* mutations into the double deletion background and observed that centromere pairing was rescued (Fig. 5e). In addition, an extra copy of *mad2* (*mad2^OE^*) in the double deletion strain, but not *mad2^OE^* alone, led to a >2-fold increase in meiotic unpairing in late pachytene oocytes (Fig. 5e). These data indicate that Mad2 negatively regulates centromere pairing when the satellite DNA arrays are disrupted. Taken together, our data suggest that Pch2 plays an important role in centromere pairing, with Mad2 inactivation as one of its probable mechanisms.

### Centromere unpairing is strongly correlated with programmed cell death

Meiotic recombination and the formation of crossovers (COs) between homologous chromosomes are critical for accurate chromosome segregation during the first meiotic division^1^. However, COs do not always form between homologues and even when they do, they can be positioned far away from the centromeres such that the tension-dependent bipolar attachment at the kinetochore may be compromised^1^ (Extended Data Fig. 8a). In these instances, centromere pairing and pericentromeric heterochromatin associations are thought to safeguard faithful chromosome segregation^16–18,33,36^. Consistently, loss of centromere pairing has been associated with increased frequency of chromosome mis-segregation during meiosis I^76,77^. To test this, we elicited centromere unpairing using germline knockdown of the SC proteins, C(2)M and C(3)G^30,31^, and observed a significant increase in X chromosome mis-segregation (non-disjunction) (Extended Data Fig. 8b).

Curiously, we did not observe X chromosome mis-segregation across many of the satellite DNA deletion genotypes, even in cases where ∼40% of late pachytene oocytes exhibit centromere unpairing (Extended Data Fig. 8c-e). These observations were reminiscent of findings from *C. elegans* where high levels of homologue unpairing only yield a subtle increase in chromosome mis-segregation^78,79^. Rather, pairing defects trigger meiotic arrest and cell death in C. elegans^69^, thus eliminating oocytes before potential mis-segregation. During *Drosophila* oogenesis, a quality control checkpoint known as mid-oogenesis cell death (MOCD) culls defective oocytes prior to the energy-intensive process of yolk production at Stage 9/10^80^ (Fig. 5f). Egg chambers with MOCD contain telltale pyknotic nuclei that are stained intensely by the DNA dye, DAPI (arrowhead, Fig. 5f). We quantified the percentage of MOCD in Stg. 7 and 8 egg chambers across all the previously described genotypes (Fig. 5g) and observed a striking correlation between meiotic unpairing and MOCD across all genotypes tested (R^2^ = 0.86, Fig. 5g, h, Extended Data Fig. 8f, g). Our data suggests that MOCD culls egg chambers containing oocytes with unpaired homologues, prior to their development into mature eggs that will be prone to chromosome mis-segregation and aneuploidy. We further tested this notion in a genetic background (*mei-P22* mutant) where we can trigger chromosome mis-segregation by blocking recombination. As expected, we observed a significant increase in Chr. X NDJ in the absence of *mei-P22* (Extended Data Fig. 8h), which was associated with a slightly elevated occurrence of MOCD (Fig. 5h). Strikingly, inducing MOCD in the *mei-P22* mutant background through heterozygous *X^Δ^* and *2^Δ^* satellite DNA deletions (*X^Δ^*/+; *2^Δ^*/+; *mei-P22^P22^*/*mei-P22^P22^*) rescued chromosome segregation fidelity (Extended Data Fig. 8h), despite elevated levels of centromere unpairing (Fig. 5g, Extended Data Fig. 5c). In contrast, homozygosity for *X^Δ^*, which does not induce MOCD (Fig. 5g), did not rescue chromosome segregation fidelity when introduced into the *mei-P22* mutant background (*X^Δ^*/ *X^Δ^*; *mei-P22^P22^*/*mei-P22^P22^*, Extended Data Fig. 8h). We also noted that ovaries lacking C(2)M and C(3)G, which produce aneuploid gametes (Extended Data Fig. 8b) exhibited low rates of MOCD (3.33%, n=360 and 3.03%, n=363) that were similar to the undepleted control (5.4%, n=241). Therefore, although the exact mechanism that triggers MOCD remains unclear, our data clearly indicate that this quality control checkpoint is a powerful culling mechanism that prevents the maturation of aneuploidy-prone oocytes. Moreover, females exhibiting higher levels of MOCD (∼20%) exhibited reduced fertility (Extended Data Fig. 8i, j), highlighting a link between compromised meiotic pairing and organismal fitness.

## Discussion

The recognition of homologous chromosomes is paramount to the success of meiosis and sexual reproduction. In this study, we have demonstrated that non-coding satellite DNA repeats function to discriminate between homologous and non-homologous chromosomes in *Drosophila* meiosis, providing experimental evidence for a hypothesis that was raised nearly 50 years ago^15^. We propose that it is the accumulated strength of multiple interactions between identical satellite DNA repeats on homologous chromosomes that ensures robust meiotic pairing. These pairing interactions likely involve sequence-specific repeat-binding proteins (e.g. D1) that mediate inter-chromosomal satellite DNA associations^55,81^ but could also occur directly through non-duplex DNA contacts^82^ or utilize non-coding RNA as intermediates^6^. The fact that these chromosome-specific satellite DNA ‘barcodes’ occur in a wide variety of eukaryotes, including primates^14,83,84^, suggests that repeat homology could be a widespread mechanism facilitating chromosome recognition.

Interestingly, the pairing defects that we observe in strains carrying satellite DNA mismatches are largely limited to the centromeric and pericentromeric heterochromatin, while pairing and synapsis at the chromosome arms appear to be unaffected. These observations suggest that satellite DNA barcodes contribute to ‘local’ homologue recognition, similar to other previously described pairing mechanisms^6^. This repeat-based homology detection may be supported by other pairing mechanisms (e.g. recombination) and loci present elsewhere^43–45^ to ensure accurate pairing along the entire chromosome. For example, in *Drosophila*, SC polymerization is initiated from the centromeric and pericentromeric heterochromatin and subsequently at the chromosome arms, likely at CO sites^30,31^. Since the centromeric and pericentromeric heterochromatin tend to exhibit fewer meiotic double strand breaks^85^ and suppress recombination^86^, we consider it likely that repeat-based homology detection is likely critical for accurate partner choice at these regions, especially since the SC does not directly monitor sequence homology.

Consistently, our data indicates that satellite DNA barcodes are important during the establishment of meiotic pairing in *Drosophila* (region 1 of the germarium)^34,35^. Therefore, we propose that recombination-based and repeat-based pairing are partially redundant mechanisms, which are required to elicit a reliable outcome from occasionally unreliable parts.

Our findings on satellite DNA function in meiotic pairing align with observations made in the nematode *C. elegans*, where chromosome-specific repetitive loci known as pairing centers (PCs), along with their associated binding proteins, facilitate homologue pairing^78,79,87–90^. Heterozygosity for PCs disrupts pairing across meiotic prophase and leads to apoptosis-mediated elimination of unpaired oocytes. These findings suggest that organisms such as *C. elegans* and *D. melanogaster*, whose chromosomes can pair and synapse without recombination^2,3^, have evolved parallel strategies to safeguard meiotic fidelity. Since the human genome also contains chromosome-specific satellite DNA ‘barcodes’^14,84^, we suggest that similar repeat-based pairing mechanisms could be an important safeguard in human meiosis and protect against anueploidy.

Lastly, satellite DNA are known to evolve extremely rapidly in eukaryotes, mutating at rates that are around 3-6 orders of magnitude higher than non-repetitive sequences^91^. However, the satellite DNA content across individuals of a species tends to be relatively uniform, especially in comparison to the dramatic differences in the satellite DNA content between species^12,28,61,92^. We propose that the role of satellite DNA in homologue pairing that we have identified in this study potentially constrains satellite DNA evolution. Since meiotic progression has a direct effect on fitness, natural selection may favour the maintenance of similar satellite DNA arrays on homologous chromosomes within interbreeding populations. At the same time, geographical isolation of individuals may allow for gradual changes in satellite DNA content, potentially reducing reproductive compatibility with their progenitors and paving the way for speciation.

## Methods

### Fly husbandry and strains

All fly stocks were raised on standard Bloomington medium at 25°C unless otherwise indicated. *D. melanogaster y w* was used as a wild type stock. *Zhr^1^* (BDSC25140), *Df(2R)^M41A10^* (BDSC741), *Df(2R)^Nip-D^(BDSC8475)*, *Df(2L)^c’^*(BDSC4959), *T(1;2)^l-v25^* (BDSC3817), *mCherry^RNAi^* (BDSC35785), *C(2)M^RNAi^* (BDSC43977), *C(3)G^RNAi^* (BDSC62969), *pch2^EY01788a^* (BDSC82769), *Df(3R)^Bsc195^* (BDSC9621), *mad2^EY21687^* (BDSC22495), *mei-P22^P22^* (BDSC4931) and *C(1)DX, y^1^f^1^Bx^0^/y^1^w^a^/Dp(1;Y)Bar^S^* (BDSC4300) were obtained from the Bloomington Drosophila Stock Center. *D. simulans Lhr^1^* and *In(1)AB* were obtained from Daniel Barbash^63^. *mad2^P^* and *GFP::Mad2* were gifts from Régis Giet and Roger Karess^93,94^. *nos-GAL4^+VP16^* has been previously described^95^ and *nos-GAL4^+VP16^* on Chr. 2 was a gift from Yukiko Yamashita. The Global Diversity Lines^60^ were obtained from Roman Arguello. The *D1^KO^* CRISPR allelle has been recently described^56^.

### Fertility and segregation assays

For female fertility assays, a single tester female was maintained with two *y w* males in vials for 7d and the number of resulting progeny was counted until 20 days post setup. For the segregation assays, a single tester female was maintained with two *y^1^w^a^/ Dp(1;Y)Bar^S^*males in vials for 7d and the number of progeny were counted until 20 days post setup. Progeny arising from X chromosome non-disjunction were determined based on the dominant Bar-of-stone marker mutation on the Y chromosome. Vials that contained any deceased parent flies were omitted from the analysis.

### Immunofluorescence staining

For dCenp-A staining, ovaries of 4-day-old adult females were dissected in 1x PBS, the ovarioles were partly separated using fine tweezers and transferred to 4% EM-grade paraformaldehyde in PBS and fixed for 30 minutes at room temperature on a nutator. For immunostaining, fixed tissues were washed in 1x PBS-T (PBS containing 0.1% Triton-X) three times for 15 min each, followed by incubation with primary antibody in 3% bovine serum albumin (BSA) in PBS-T at 4°C overnight on a nutator. On day 2, samples were washed as above, incubated with secondary antibody in 3% BSA in PBS-T at 4°C overnight, washed as above, and mounted in VectaShield with DAPI (Vector Labs). anti-dCenp-A (gift from Christian Lehner) was used at a final concentration of 1:1000. For immunostaining of the synaptonemal complex, ovaries of 4-day-old adult females were dissected in 1x PBS, the ovarioles were partly separated using fine tweezers and transferred to the fixative solution [2% Formaldehyde, 0.5% *Pierce NP40* and heptane] and fixed for 20 minutes at room temperature on a nutator. The tissues were washed, stained and imaged as mentioned above. The monoclonal mouse anti-C(3)G 1A8-1G2 antibody^96^ was used at a final concentration of 1:1000. The stages of oogenesis were estimated based upon a previous study^97^. Mid-oogenesis cell death (MOCD) was quantified as the percentage of dying egg chambers in Stg. 7 and 8. Fluorescent images were taken using a Leica TCS SP8 confocal microscope with 63x oil-immersion objectives (NA=1.4). Images were processed using Adobe Photoshop or ImageJ.

### Oligopaint DNA FISH

Three pairs of *Drosophila ovaries* were dissected and fixed as described above. The tissue was the washed in 2× SSC-T with increasing formamide concentrations (20%, 40%, and 50%) for 10 min each followed by a final 10 min wash in 50% formamide. Next, samples in 50% formamide plus 2× SSC-T were transferred to a PCR tube and incubated for 4 h at 37°C, 3 min at 92°C, and 20 min at 60°C. After this step, excess formamide solution was removed, and the hybridization mix (20–40 pmol per probe; 36 µL of probe buffer plus 1 µL of RNase A) was added to the ovaries. Samples were denatured for 3 min at 91°C followed by overnight incubation at 37°C in the dark. Following hybridization, samples were first rinsed with 50% formamide plus 2× SSC-T and then washed twice for 30 min each at 37°C. Next, samples were washed once with 20% formamide plus 2× SSC-T for 10 min at room temperature, washed four times with 2× SSC-T for 3 min each, and then mounted with VectaShield plus DAPI. The probes against Locus A, B and C have been previously described^98^.

### Satellite DNA FISH

Whole mount *Drosophila* ovaries were prepared as described above, and optional immunofluorescence staining protocol was carried out first. Subsequently, samples were postfixed with 4% formaldehyde for 10min and washed in PBS-T for 30min. Fixed samples were incubated with 2mg/ml RNase A solution at 37 °C for 10min, then washed with PBS-T+1mM EDTA. FISH using heat denaturation was carried out as follows: samples were washed in 2xSSC-T (2xSSC containing 0.1% Tween-20) with increasing formamide concentrations (20%, 40%, and 50%) for 15min each followed by a final 30-min wash in 50% formamide. Hybridization buffer (50% formamide, 10% dextran sulfate, 2× SSC, 1mM EDTA, 1μM probe) was added to washed samples. Samples were denatured at 91 °C for 2min, then incubated overnight at RT. For mitotic chromosome spreads, larval 3^rd^ instar brains were squashed according to previously described methods^99^. Briefly, tissue was dissected into 0.5% sodium citrate for 5–10min and fixed in 45% acetic acid/2.2% formaldehyde for 4–5min. Fixed tissues were firmly squashed with a cover slip and slides were submerged in liquid nitrogen until bubbling ceased. Coverslips were then removed with a razor blade and slides were dehydrated in 100% ethanol for at least 5min. After drying, hybridization mix (50% formamide, 2× SSC, 10% dextran sulfate, 100ng of each probe) was applied directly to the slide, samples were heat denatured at 95 °C for 5min and allowed to hybridize overnight at room temperature. Following hybridization, slides were washed three times for 15min in 0.2× SSC and mounted with VectaShield with DAPI (Vector Labs). The following satellite DNA probes were used: Rsp – 5’ ATTO488-GGA AAA TCA CCC ATT TTG ATC GC 3’, Dodeca – 5’ Cy5-ACC GAG TAC GGG ACC GAG TAC GGG ACC GAG TAC GGG 3’, AAGAC – 5’ Cy5-AAG ACA AGA CAA GAC AAG ACA AGA CAA GAC 3’, AAGAG – 5’ Cy3-AAG AGA AGA GAA GAG AAG AGA AGA GAA GAG 3’, IGS 5’ ATTO488-AGT GAA AAA TGT TGA AAT ATT CCC ATA TTC TCT AAG TAT TAT AGA GAA AAG CCA TTT TAG TGA ATG GA 3’, 359 – 5’ Cy3-AGG ATT TAG GGA AAT TAA TTT TTG GAT CAA TTT TCG CAT TTT TTG TAA G 3’, AACAC – 5’ Cy3-AAC ACA ACA CAA CAC AAC ACA ACA CAA CAC 3’, AATAG – 5’ Cy5-AAT AGA ATA GAA TAG AAT AGA ATA GAA TAG-Cy5 3’ and Prod – 5’ Cy5-AAT AAC ATA GAA TAA CAT AGA ATA ACA TAG 3’. The probe for the 260bp satellite DNA has been previously described^28^. To estimate satellite DNA abundance in the GDL strains, we analyzed mitotic chromsomes from either single slices or composite images from a maximum projection of 2-3 slices in the progeny of selected strains using Fiji (ImageJ). First, we measured cumulative intensity of the AAGAG satellite DNA on individual chromosomes. The homologue with the lower AAGAG intensity was designated the I16 chromosome while the other homologue was designated as the chromosome from the other GDL strain i.e. ZH33 or B59. Based on these designations, the cumulative intensity of other satellite DNA repeats were measured on their respective chromosomes and quantified as an intensity ratio. This method can detect satellite DNA repeats with a minimal abundance of approximately 30-100kb^28^.

### Electrophoretic mobility shift assay

The radioactively labelled dsDNA substrates were prepared by amplifying the control (50%GC 260bp and 359bp) and satellite DNA (260bp and 359bp repeat monomer) sequences from plasmid substrates using a PCR reaction containing 66 nM [α-32P]dCTP with the standard dNTPs concentration (200 µM each). The purification of mNeonGreen (mNG)-tagged D1 protein has been recently described^56^. The reactions (15 µl volume) were performed in binding buffer containing 25 mM Tris-HCl pH 7.5, 1 mM DTT, 3 mM MgCl_2_, 0.1 mg/ml BSA (New England Biolabs), 0.3 nM radioactively labelled PCR-based dsDNA substrate and 75 ng of linearized plasmid competitor. After the addition of D1 protein, the reactions were incubated on ice for 15 min. Loading dye (50% glycerol, bromophenol blue) was added and the products were separated by 0.6% agarose gel electrophoresis in Tris-Acetate-EDTA (TAE) buffer. The gels were dried on a DE81 paper (Whatman) and exposed to storage phosphor screen (GE Healthcare) and scanned by a Typhoon Phosphor Imager (FLA 9500, GE Healthcare).

### Bead clustering assay

For the bead clustering assay, we adapted a previously described protocol. Briefly, we functionalized 6.60µl of streptavidin-coated magnetic beads (Dynabeads™ M-280, 10mg/ml) with 28.14ng of biotinylated DNA (359bp satellite DNA monomer, 260bp satellite DNA monomer and 359bp 50%GC control). 30 µL reactions were assembled in clear 96-well plates at room temperature in BW (10 mM Tris-HCl pH 7.5, 5 mM MgCl₂). Each well contained 0.3nM DNA on beads (0.009 pmol DNA and ∼0.005 mg beads), 3µg/ml of unlabeled plasmid competitor and purified mNG-D1 at 0, 40, 50, or 60 nM. Plates were placed on a magnetic rack to aggregate the paramagnetic beads, briefly vortexed to resuspend, and then incubated for 20 min at room temperature prior to imaging. Bright-field images were acquired on a Leica TCS SP8 confocal microscope with 20x objective with fixed exposure and magnification across conditions. Images were analyzed in Fiji (ImageJ). For each field, a global threshold was set manually to segment beads; obvious artifacts (e.g., threads/fibers, air bubbles) were manually removed from the binary mask/ROI Manager. Particle analysis was performed with size and shape filters (size ≥ 3–5 µm²; circularity 0.15–1.00). For each condition, particle size distributions (area, Feret’s diameter) and summary statistics were quantified from three replicates.

### RNA sequencing and data analysis

For each sample, approximately 5 pairs of ovaries from 0-1 day old females were dissected in RNase-free 1x PBS and flash frozen in liquid nitrogen until RNA extraction. RNA extraction for each replicate was performed using the RNeasy RNA extraction kit (Qiagen). Samples were treated with DNase post RNA extraction and purified using an RNA purification kit (Promega). RNA concentrations were assessed using a Nanodrop as well as a Qubit RNA analyzer for sample quality and RIN scores. rRNA-depleted libraries were prepared and sequenced by the Functional Genomics Center Zurich (FGCZ) on an NovaSeq 6000 (Illumina) using paired-end 100 bp sequencing. The resulting raw reads were cleaned by removing adaptor sequences, low-quality-end trimming, and removal of low-quality reads using BBTools v 38.18 (Bushnell, B. BBMap. Available from: https://sourceforge.net/projects/bbmap/). The exact commands used for quality control can be found on the Methods in Microbiomics webpage^100^. The qualitycontrolled reads were aligned against BDGP6.32 using STAR aligner^101^. Differential gene expression analysis was performed using Bioconductor R package DESeq2 v1.37.4^102^.

### Single nucleotide polymorphism and deletions datasets

The SNP, satellite DNA and indel datasets were obtained from previous studies on the GDL strains^60,61^. Briefly, SNPs were limited to autosomes (excluding chromosome 4) and only small intronic (positions 32–65bp) and 4-fold degenerate positions were used based on genomic annotations generated using SNPeff (167857 SNPs). The SNP data were further filtered using vcftools v0.1.16^103^ with the following parameters: --remove-indels --mac 1 --minQ 30 --min-meanDP 5 --max-meanDP 100 --minDP 5 --maxDP 100 --max-missing 0.9 and pruned to remove linked SNPs using plink2 v2.0.0^104^ with the following parameters --indep-pairwise 50 10 0.2. The filtered file contained 50034 final SNPs. The indel dataset was filtered using vcftools v0.1.16 with the following parameters --minQ 30 --minDP 5 --maxDP 100 --min-meanDP 5 --max-meanDP 100 --maf 0.05 --max-missing 0.9. Multiallelic variants were split into separate biallelic records using bcftools v.1.21^105^ with norm -m parameters. Deletions/insertions on Chr X were excluded from the analysis. The final filtered file contained 66385 insertions/deletions.

### Phylogenetic tree construction and population structure analysis

The filtered SNP dataset was used to generated ML tree using iqtree v2.4.0^106^ with the following parameters -m MFP+ASC, with SYM+ASC+R6 model selected as best fit.The final tree ML was visualized using iTol^107^. To investigate the population structure of GDL lines, we performed principal component analysis on the SNP and indel datasets using plink2 v2.0.0^104^.

## Data Availability Statement

The RNA-seq datasets will be uploaded to a publicly accessible repository prior to publication.

## Acknowledgements

We thank members of the Jagannathan lab, Hugo Stocker, Yukiko Yamashita, Scott Hawley and Joao Matos for discussion and comments on the manuscript. We are grateful for the reagents and resources provided by Scott Hawley, Christian Lehner, Eric Joyce, Son Nguyen, Sharon Bickel, the Bloomington Drosophila Stock Center, the Vienna Drosophila Resource Center, the Kyoto Drosophila Stock Center and the Developmental Studies Hybridoma Bank. We thank Andrew Clark and Roman Arguello for sequencing data related to the GDL strains. We thank Katarzyna Nowak, Melanie Pfister and Lilly Schneider for assistance with the fertility and segregation assays. We acknowledge microscopy support from the Scientific Center for optical and Electron Microscopy (ScopeM) and NGS support from the Functional Genomics Center Zurich (FGCZ). We also thank ETH Zurich IT services and High Performance Computing facilities for computational support. M.J. is funded by the Swiss National Science Foundation (310030_189131 and 320030_228043). P.C. is funded by the Swiss National Science Foundation (310030_207588 and 310030_205199) and the European Research Council (101018257). S.S. acknowledges core funding from ETH Zürich.

## Author Contributions

LS performed most of the experiments. Other experiments were done by AC, IC and CJR. AS performed all bioinformatic analysis. MJ conceived and supervised the project; MJ and LS wrote the manuscript with input from all authors.

## Competing Interests

The authors declare no competing interests

## Notes

### Competing Interest Statement

The authors have declared no competing interest.

