## Supplementary Figures 1-8 for "Meiotic pairing through barcode-like satellite DNA repeats"

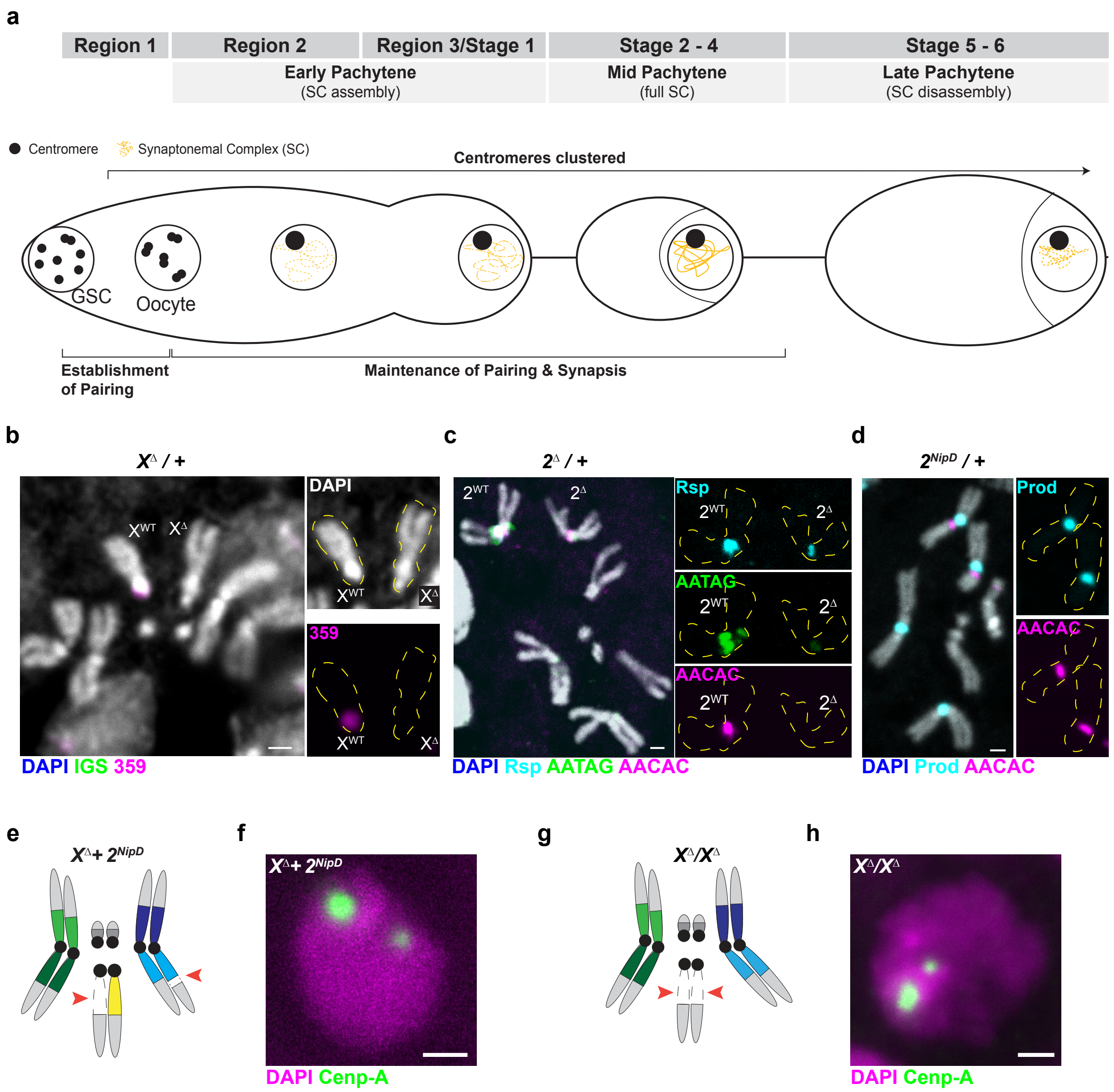


**Extended Data Fig. 1. *Drosophila* oogenesis is a powerful model to study meiotic pairing. a,** Schematic of the dynamics of homologue pairing and synapsis across *Drosophila* oogenesis **b,** FISH against the 359bp satellite repeat (magenta) and co-stained with DAPI (white) on mitotic chromosomes from a strain heterozygous for $X^{\Delta}$ (*Zhr*$^{1}$). **c,** FISH against the $(AATAG)_n$ satellite repeat (green), the $(AACAC)_n$ satellite repeat (magenta) and the Rsp satellite repeat (cyan) and co-stained with DAPI (white) on mitotic chromosomes from a strain heterozygous for $2^{\Delta}$ (*Df(2R)*$^{M41A10}$). **d,** FISH against the $(AACAC)_n$ satellite repeat (magenta) and the Prod/$(AATAACATAG)_n$ satellite repeat (cyan) and co-stained with DAPI (white) on mitotic chromosomes from a strain heterozygous for $2^{NipD}$ (*Df(2R)*$^{Nip\text{-}D}$). **e-h,** A schematic of the karyotype and IF against dCenp-A (green) in late pachytene oocytes co-stained with DAPI (magenta) from the indicated genotypes. Red arrows indicate satellite DNA and chromosomal deletions. Scale bar, 1μm.

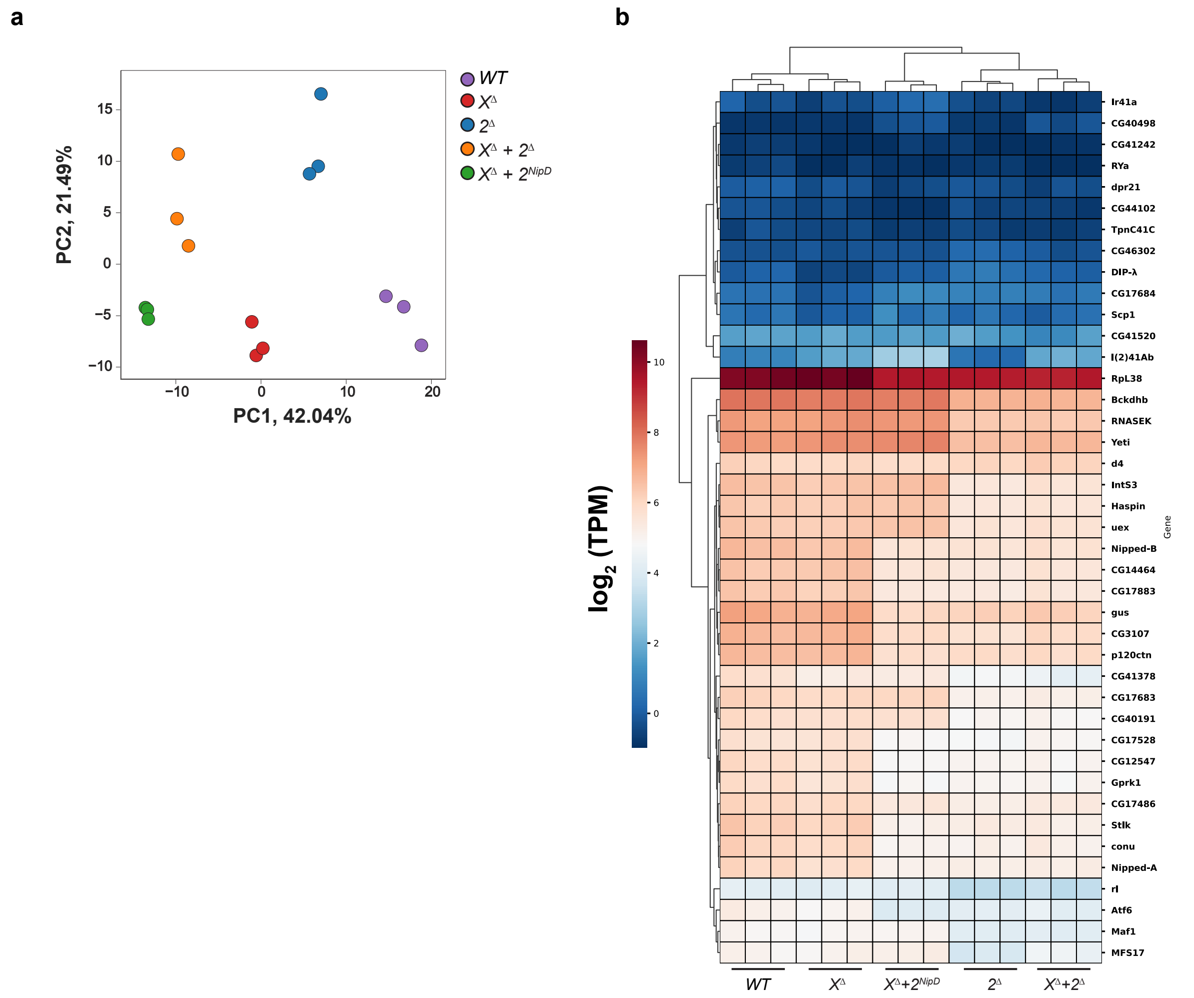


**Extended Data Fig. 2. Estimating of gene expression changes following satellite DNA deletion. a,** Principal component analysis (PCA) of gene expression from 0-1d old ovaries from the indicated genotypes. **b**, Expression of the indicated genes ($log_2$TPM) across the indicated genotypes.

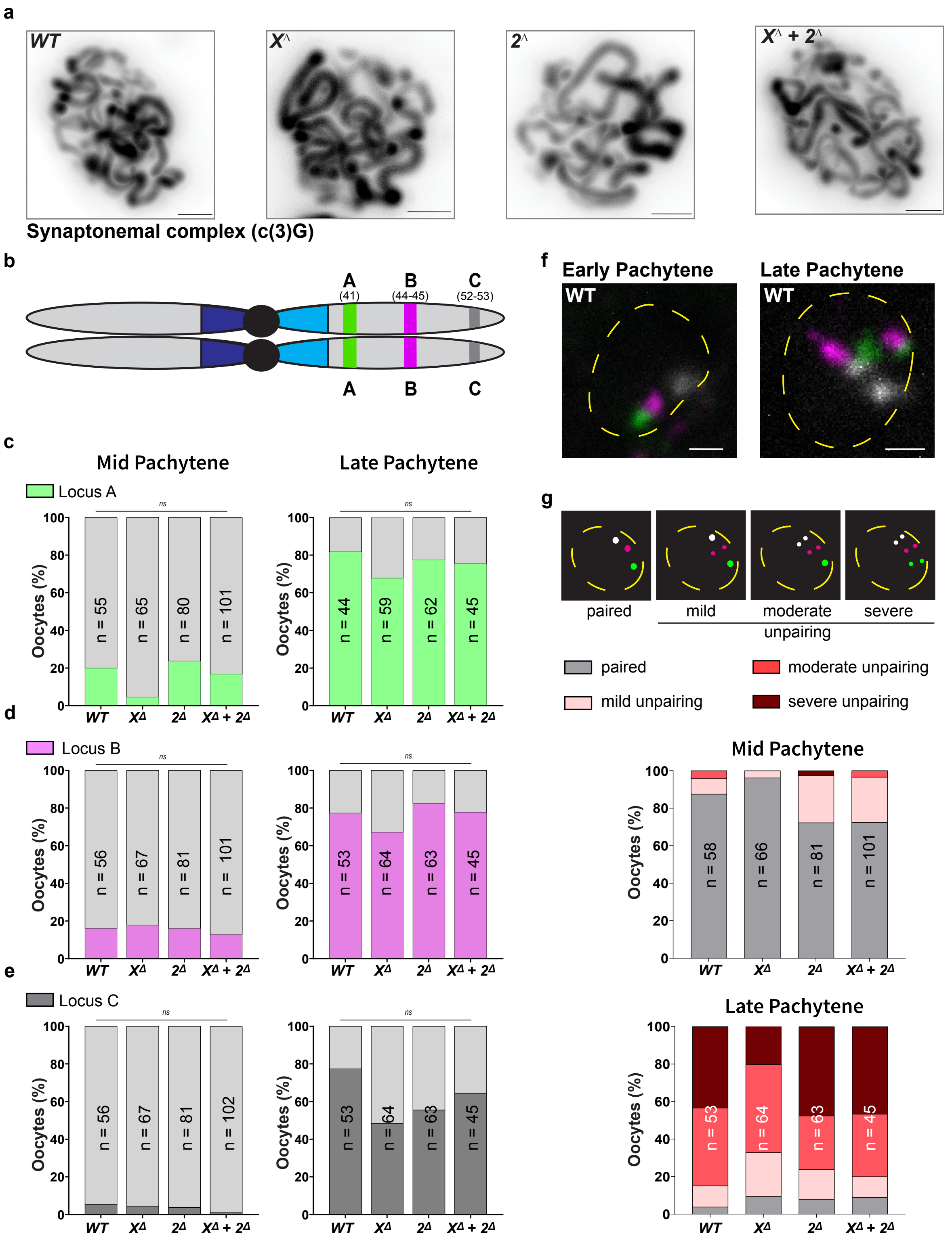


**Extended Data Fig. 3. Chr. 2R pairing and synapsis is unaffected by satellite DNA deletions. a,** IF against the transverse filament of the synaptonemal complex C(3)G (black) in mid pachytene oocytes from the indicated genotypes. **b,** Schematic of Chr. 2 with the location of three 1Mb loci (A (green), B (magenta) and C (white)) depicted on Chr. 2R**. c-e,**

Quantification of mid  and late pachytene oocytes with unpaired locus A **(c)**, locus B **(d)** and locus C **(e)** from the indicated genotypes. **f,** Oligopaint-based DNA FISH against Chr. 2R-A (green), Chr. 2R-B (magenta) and Chr. 2R-C (white) in mid  and late pachytene oocytes from a *WT* strain. Dashed line marks the nuclear boundary. **g,** (top) Schematic showing oocytes mild, moderate or severe unpairing at Chr. 2R. (bottom) Quantification of oocytes with mild, moderate or severe unpairing of Chr. 2R in mid  and late pachytene oocytes from the indicated genotypes. *n* indicates the number of oocytes analyzed and ns indicates $p > 0.05$ from a Fisher's exact test.  All scale bars are 1μm.

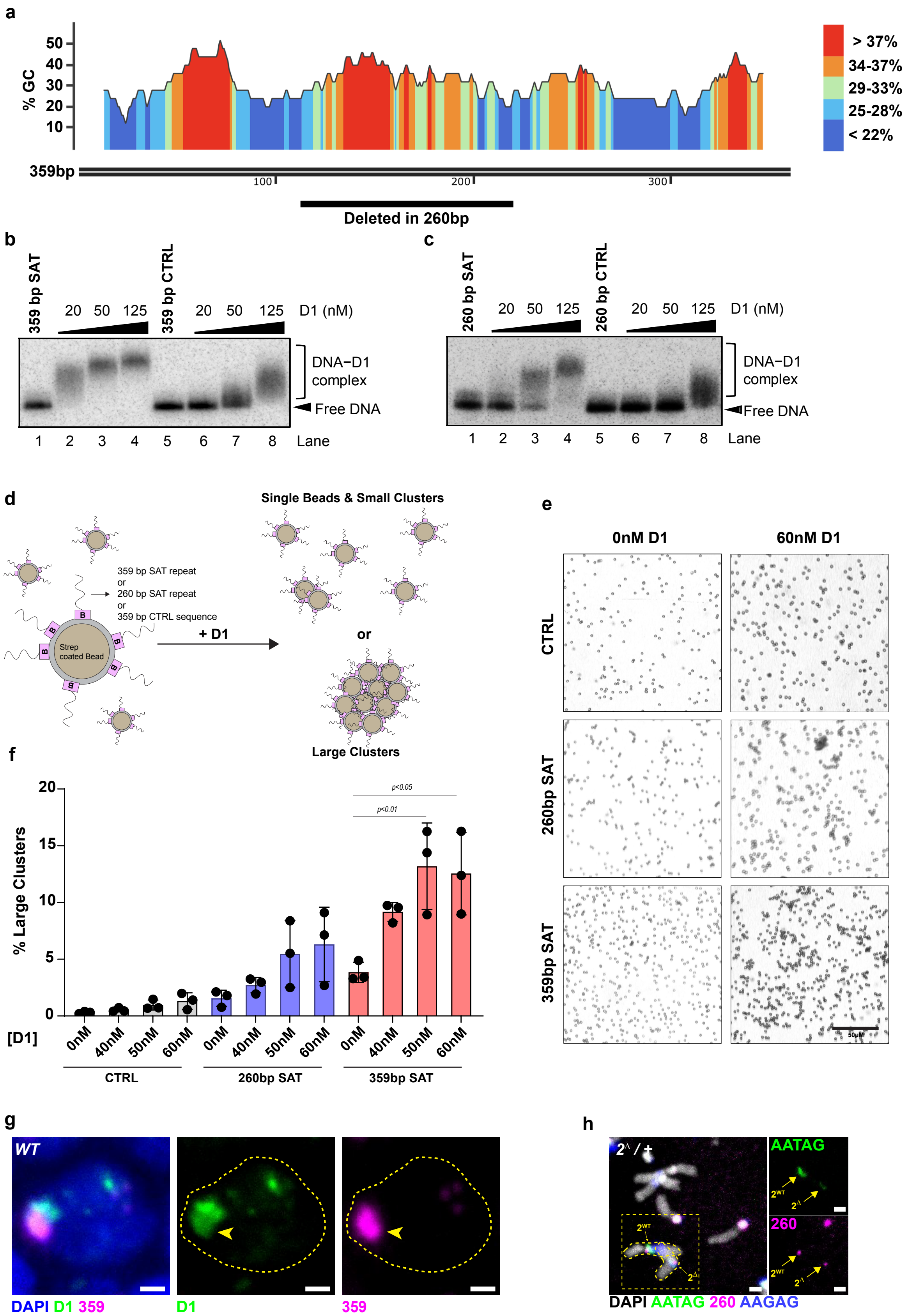


**Extended Data Fig. 4. The satellite DNA-binding protein D1 binds and clusters the 359bp and 260bp satellite DNA repeats in vitro. a,** %GC content of the 359bp satellite

DNA repeat. Black line indicates the ~100bp region deleted in the 260bp variant. **b, c,** Electrophoretic mobility shift assay (EMSA) of increasing concentrations of purified D1 with the 359bp satellite DNA monomer (**b**) or the 260bp satellite DNA monomer (**c**) with 50% GC length-matched controls. **d,** Schematic of the bead clustering assay. Streptavidin-coated beads were coupled to the following biotinylated dsDNA – 359 bp satellite DNA monomer (359bp SAT), 260bp satellite DNA monomer (260bp SAT) or a 359bp 50% GC control (CTRL) – and incubated with purified D1 in the presence of an unlabelled plasmid DNA competitor. The assay detects varying degrees of bead clustering, ranging from dispersed beads to small or large clusters. **e,** Brightfield images showing bead clusters from the indicated conditions. Scale bar: 50 µm. Image analysis was used to quantify individual beads, small bead clusters (3-6 beads) and large bead clusters. **f,** Quantification of percentage of large bead clusters (>3-6 beads) in the indicated conditions following thresholding and automated particle size analysis (see methods). *p* values were obtained using one-way ANOVA followed by Dunnett's multiple comparisons test. **g,** FISH against the 359 bp satellite DNA (magenta) in a pre-meiotic female germ cell stained for D1 (green) and DAPI (blue). Dashed line marks the nuclear boundary and arrowheads indicate co-localization between D1 and the 359bp satellite DNA repeat. **h,** FISH against the $(AAGAG)_n$ satellite repeat (blue), $(AATAG)_n$ satellite repeat (green), 260bp satellite repeat (magenta) co-stained with DAPI (white) on mitotic chromosomes from a strain heterozygous for $2^{\Delta}$ ($Df(2R)^{M41A10}$). Scale bar: 1 µm.

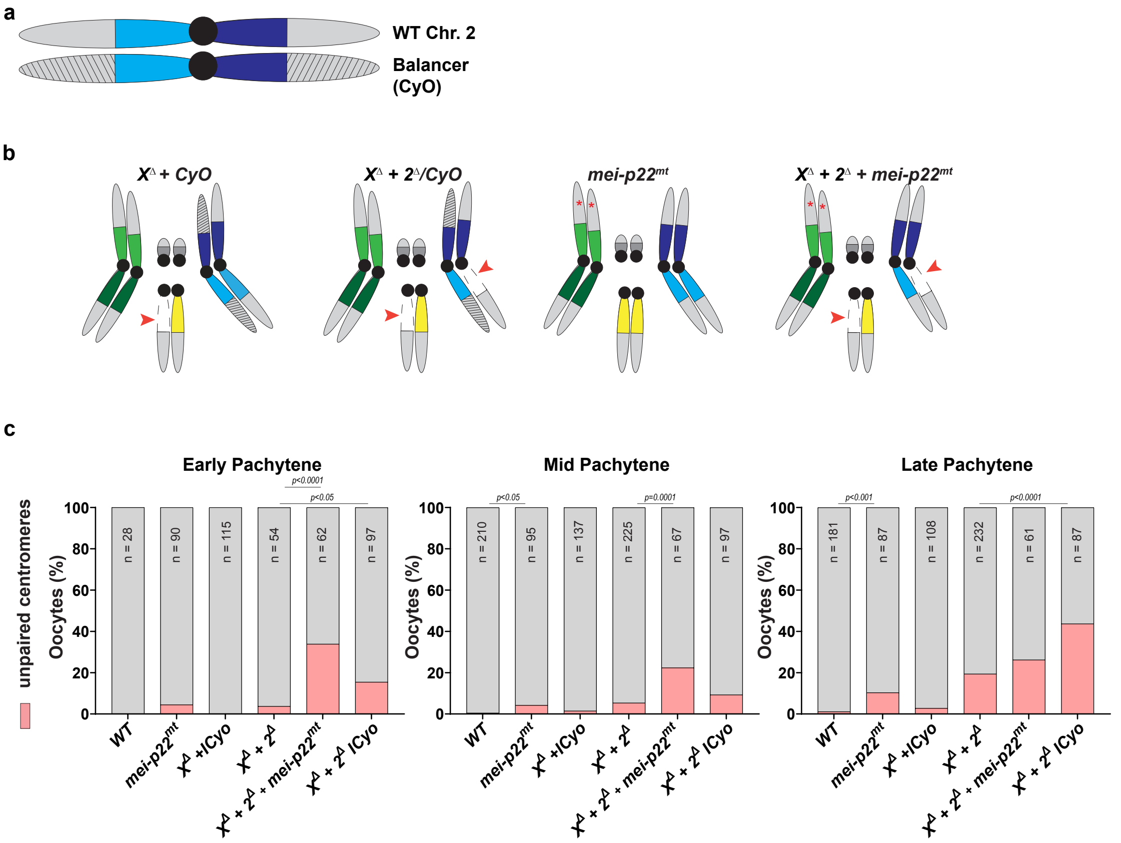


**Extended Data Fig. 5. Arm synapsis and recombination suppresses centromere unpairing following satellite DNA deletion. a,** Schematic of a strain heterozygous for a Chr. 2 balancer (CyO) containing multiple paracentric inversions (gray lines). **b,** Schematic of the genotypes used in this figure. Red arrows indicate satellite DNA deletions. Red asterisks represent the *mei-P22*$^{P22}$ mutation. **c,** Quantification of oocytes with >4 dCenp-A foci, indicating unpaired centromeres, in early , mid  and late pachytene in the indicated genotypes. *n* indicates the number of oocytes analyzed and *P* values were obtained from a Fisher's exact test. WT and $X^{\Delta}$ + $2^{\Delta}$ unpaired centromere values were reused from **Fig. 1e**.

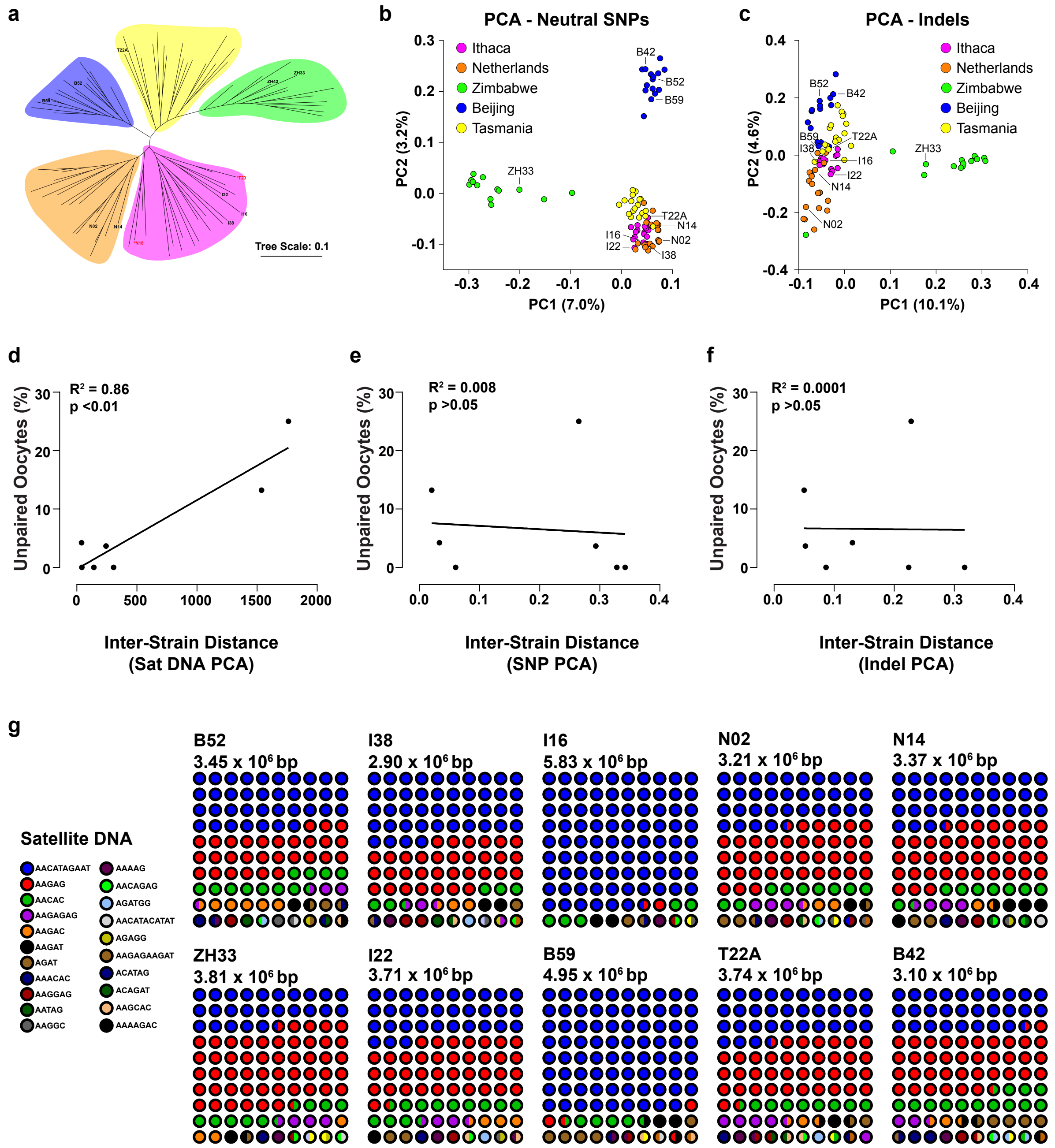


**Extended Data Fig. 6. Satellite DNA divergence is strongly correlated with meiotic pairing defects in the progeny of the GDL strains. a,** Unrooted phylogenetic tree of GDL strains constructed from neutral SNPs. Branch colors indicate geographical origin. GDL strains used in this study are shown in black, and outlier strains (N18 and T23) that cluster outside their expected geographical group are highlighted in red**. b, c,** Principal component analysis of the GDL strains based on neutral SNPs (b) and indels (c). GDL strains used in this study are labeled. **d-f,** Correlation between % oocytes with centromere unpairing in late pachytene and inter-GDL euclidean distance from the Satellite DNA PCA (**d**), neutral SNP PCA (**e**) and indel PCA (**f**). $R^2$ and *p* value were determined from a linear regression analysis. **g,** Satellite DNA composition of the indicated GDL strains along with the estimated total satellite DNA content in bp.

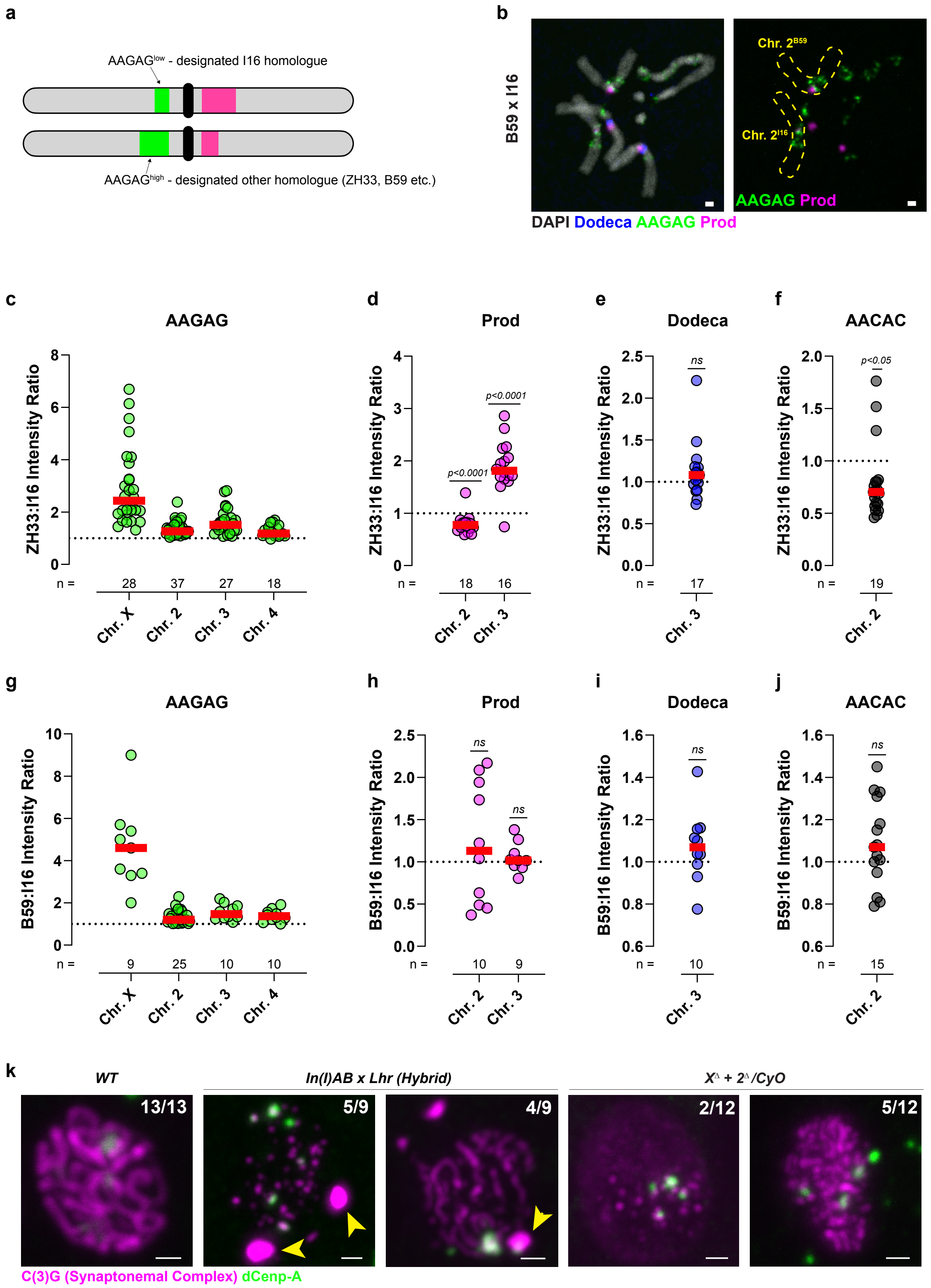


**Extended Data Fig. 7. The Chr. 2 satellite DNA barcode is substantially altered in select GDL strain pairs. a,** Schematic illustrating how relative satellite DNA abundance was measured in mitotic chromosome spreads from GDL strain crosses. See methods for details. **b,** FISH against the Dodeca satellite repeat (blue), $(AAGAG)_n$ satellite repeat (green), Prod/$(AATAACATAG)_n$ satellite repeat (magenta) co-stained with DAPI (white) on mitotic chromosomes in the progeny of B59 and I16 strains. Scale bar: 1 µm. **c - j,** Graphs showing the chromosome-specific intensity ratios of the indicated satellite DNA repeats from the designated ZH33 chromosome and the designated I16 chromosome (**c-f**) and from the designated B59 chromosome and the designated I16 chromosome (**g-j**). *n* indicates the number of chromosomes analyzed and *p* values from a one-sample *t*-test indicate a significant difference from an intensity ratio of 1. **k,** IF against the transverse filament of the synaptonemal complex C(3)G (magenta) and dCenp-A (green) in mid pachytene oocytes from the indicated genotypes. Arrowheads indicate polycomplex-like aggregates and numbers indicate the fraction of oocytes with the indicated phenotype. All scale bars are 1 µm.

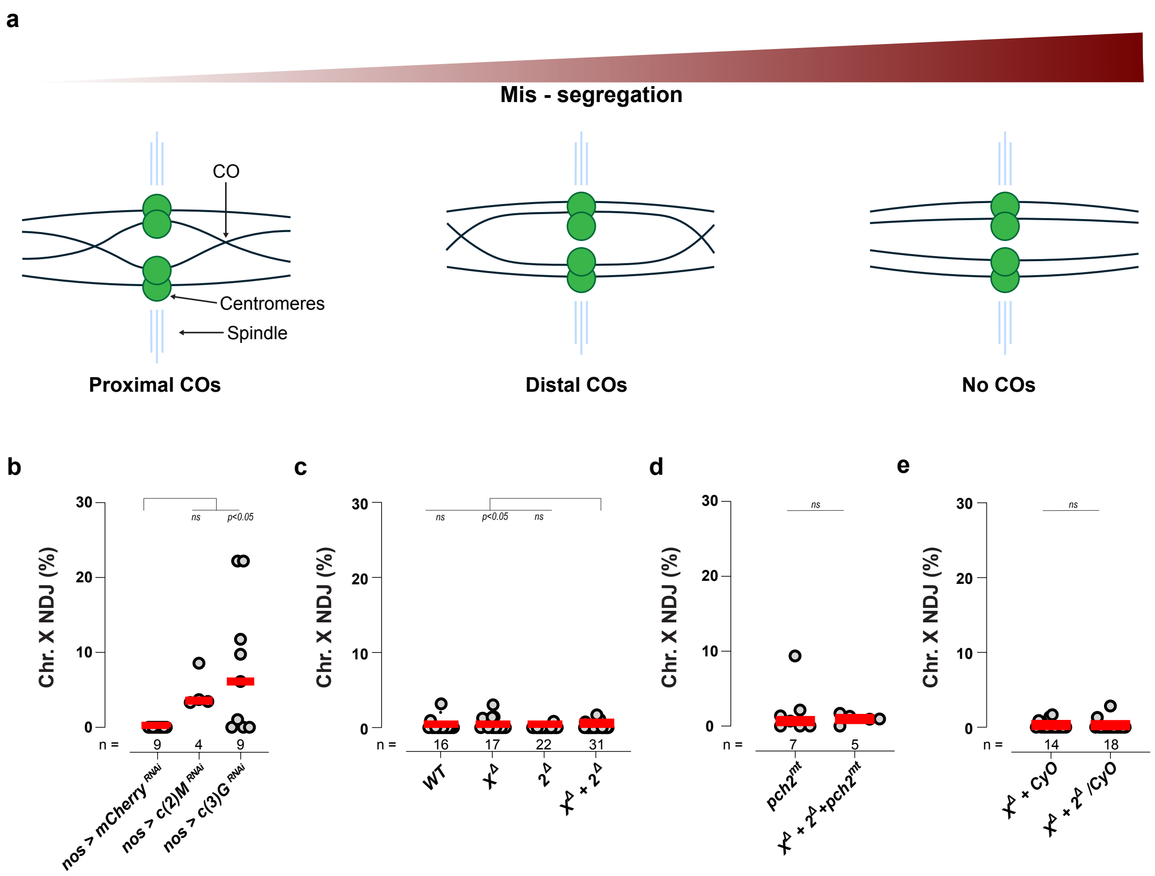


f

| Genotype | Unpaired oocytes (%) | MOCD (%) |
|---|---|---|
| *I38 x B 52* | 0.0 | 4.0 (n= 529) |
| *ZH33 x B42* | 0.0 | 4.4 (n= 437) |
| *I22 x T22A* | 0.0 | 6.3 (n= 632) |
| *In(1)AB x yw* | 1.6 | 7.8 (n= 360) |
| *I16 x B59* | 3.7 | 7.9 (n= 291) |
| *N 14 x N02* | 4.2 | 5.2 (n= 442) |
| *I16 x I 22* | 13.2 | 10.0 (n= 300) |
| *ZH33 x I16* | 25.0 | 12.2 (n= 180) |
| *In(1)AB x Lhr*$^{1}$ | 50.6 | 51.2 (n= 166) |

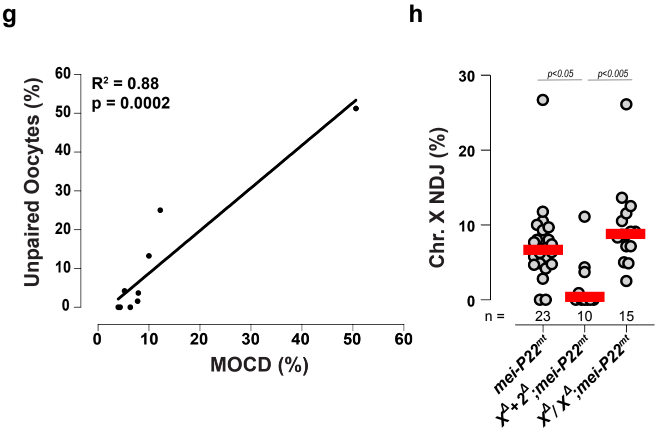


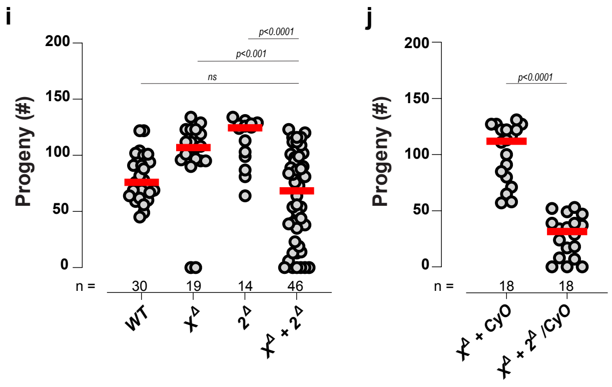


**Extended Data Fig. 8. Loss of meiotic centromere pairing does not lead to X chromosome non-disjunction. a,** Schematic of the effect of crossover (CO) position on chromosome non-disjunction during meiosis I. **b-e**, Percentage of X chromosome non-disjunction (NDJ) from the indicated genotypes. **f,** Table indicating percentage of oocytes with unpaired centromeres in late pachytene and corresponding percentage of Stg 7-9 egg chambers with MOCD from the indicated genotypes. n indicates total number of Stg 7-9 egg chambers scored. **g,** Correlation between centromere unpairing and MOCD from **(f)**. $R^2$ and *p* value were determined from a linear regression analysis. **h,** Percentage of X chromosome non-disjunction (NDJ) from the indicated genotypes. **i, j,** Number of progeny per female from the indicated genotypes. *p* values were obtained from Tukey's multiple comparisons test from an ordinary one-way ANOVA. ns indicates $p > 0.05$. Red line marks the median.
